# Mining metagenomes for natural product biosynthetic gene clusters: unlocking new potential with ultrafast techniques

**DOI:** 10.1101/2021.01.20.427441

**Authors:** Emiliano Pereira-Flores, Marnix Medema, Pier Luigi Buttigieg, Peter Meinicke, Frank Oliver Glöckner, Antonio Fernández-Guerra

## Abstract

Microorganisms produce an immense variety of natural products through the expression of Biosynthetic Gene Clusters (BGCs): physically clustered genes that encode the enzymes of a specialized metabolic pathway. These natural products cover a wide range of chemical classes (e.g., aminoglycosides, lantibiotics, nonribosomal peptides, oligosaccharides, polyketides, terpenes) that are highly valuable for industrial and medical applications^1^. Metagenomics, as a culture-independent approach, has greatly enhanced our ability to survey the functional potential of microorganisms and is growing in popularity for the mining of BGCs. However, to effectively exploit metagenomic data to this end, it will be crucial to more efficiently identify these genomic elements in highly complex and ever-increasing volumes of data^2^. Here, we address this challenge by developing the ultrafast Biosynthetic Gene cluster MEtagenomic eXploration toolbox (BiG-MEx). BiG-MEx rapidly identifies a broad range of BGC protein domains, assess their diversity and novelty, and predicts the abundance profile of natural product BGC classes in metagenomic data. We show the advantages of BiG-MEx compared to standard BGC-mining approaches, and use it to explore the BGC domain and class composition of samples in the TARA Oceans^3^ and Human Microbiome Project datasets^4^. In these analyses, we demonstrate BiG-MEx’s applicability to study the distribution, diversity, and ecological roles of BGCs in metagenomic data, and guide the exploration of natural products with clinical applications.

---

Metagenomics offers unique opportunities to mine natural product BGCs in diverse microbial assemblages from a wide range of environments^5–7^. However, given the complexity of microbial communities found in nature, and the limitations of current sequencing technologies, often only a very small fraction of the short-read sequence data can be assembled in contigs long enough to allow the identification of BGC classes. However, the annotation of individual protein domains of BGCs, is much more straightforward, given that these have comparable length to merged paired-end reads. There are several protein domains known to play important functions in the BGC-encoded enzymes. Specific domains or combinations thereof are commonly found in certain types of BGC classes. Accordingly, these are used for the automatic identification of BGC classes in genome sequences^8–10^ and to study the distribution and diversity of particular BGC classes in the environment^6,7,11–13^. Although there are various BGC mining tools with practical applications^14^, only the Natural Product Domain Seeker (NaPDoS)^11^ and the environmental Surveyor of Natural Product Diversity (eSNaPD^15^) are dedicated to the study of BGC domains. Both of these tools focus on nonribosomal peptides and polyketide synthases (NRPSs and PKSs, respectively), and take assembled or amplicon data as input. Currently, there is no technology available capable of efficiently exploiting raw metagenomic data to study the composition and diversity of natural product BGC classes and domains in the environment.

Capitalizing on the fact that BGC domains can be readily annotated in unassembled metagenomic data, and used to identify the different natural product BGC classes, we developed BiG-MEx. This tool generates ultrafast BGC domain annotations in short-read sequence data and applies a machine-learning approach to predict the BGC class coverage-based abundances (for simplicity, we will refer to these as BGC class abundance profiles). Additionally, the identified domain sequences are used to carry out a domain-based diversity analysis. This allows BiG-MEx both to deeply exploit metagenomic data, and to adapt to their ever-increasing volume. BiG-MEx consists of three interacting modules that are described below and illustrated in Fig. 1:

**Fig. 1.**
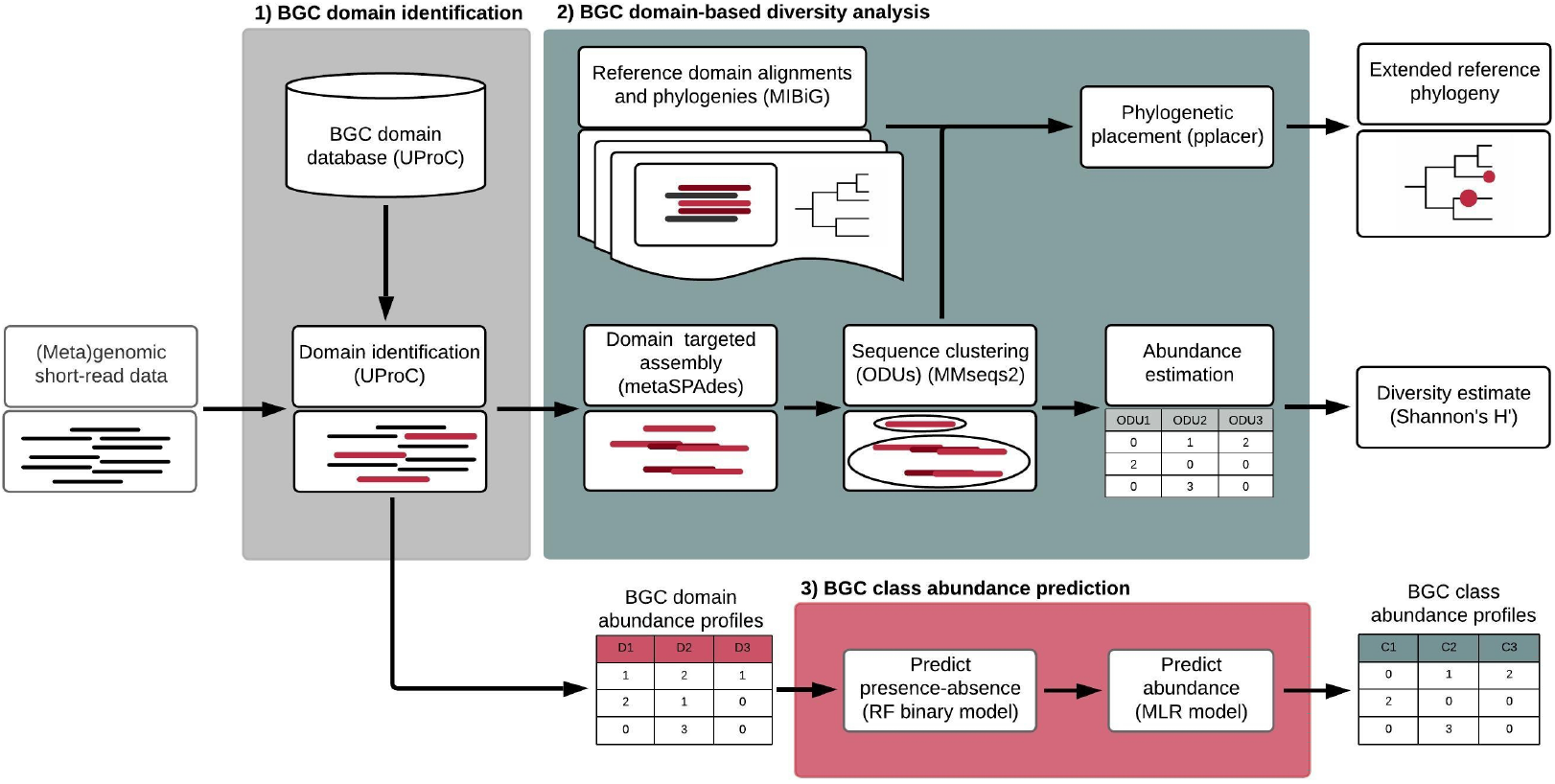
BiG-MEx analysis workflow. **1)** BGC domain identification module. To annotate the BGC domains with UProC, we created an UProC database including 150 domains, which originate from 44 different BGC classes. This database was generated based on the amino acid sequences of antiSMASH hidden Markov model (HMM) profiles^10^. Using UProC output, this module generates a count-based abundance profile of BGC domains; **2)** BGC domain-based diversity analysis module. Using the previously identified domains, this module performs a targeted assembly with metaSPAdes ^45^ to reconstruct the domain sequences. Assembled domain sequences are clustered into Operational Domain Units, and the number of ODUs and the coverage of the domain sequences within each ODU (used to approximate the abundance of the ODU) are used to compute the ODU alpha diversity. The environmental reconstructed domain sequences are placed onto reference phylogenetic trees with pplacer^48^ (maximum likelihood criteria). In this module, we include pre-computed phylogenies for 48 domains, which are based on sequence data contained in the Minimum Information about a Biosynthetic Gene cluster (MIBiG)^51^ database, allowing us to identify the relationships of query sequences with domains from pathways of known function; **3)** BGC class abundance prediction module. The domain abundance profiles are used to predict the BGC class coverage-based abundance profiles using class-specific machine-learning models. These models consist of a two-step process: First, the presence/absence of the BGC class is predicted using a random forest (RF) classifier; Secondly, the abundance is predicted with a multiple linear regression (MLR) only if the class was previously predicted as present.

1. **BGC domain identification module**. We use the Ultrafast Protein domain Classification UProC^16^ tool to identify BGC protein domains in short-read sequence data. For this purpose, we created an UProC database, which includes 150 BGC domains covering 44 BGC classes.
2. **BGC domain-based diversity analysis**. This module performs a domain-targeted assembly, clusters the assembled domain sequences to create Operational Domain Units (ODUs)^17^ and computes the ODU alpha diversity. Further, assembled domain sequences are placed onto reference phylogenetic trees. The module includes pre-computed phylogenies for 48 BGC domains. These were selected based on domain sequences from experimentally characterized biosynthetic gene clusters with enough sequence information for phylogenetic analysis.
3. **BGC class abundance prediction module**. We created machine-learning models that predict the abundance of BGC classes based on the domain annotation. The models are class-specific and consist of a random forest (RF) classifier to predict the presence/absence of a BGC class, and a multiple linear regression (MLR) to predict its abundance. These models can be customised to target metagenomic and genomic data from different environments and taxa, respectively.

To evaluate the performance of BiG-MEx, we first assessed how the UProC-based domain identification used in BiG-MEx improves the data processing speed compared to HMMER^18^ (i.e., the traditional approach for domain annotation) for the annotation of the 150 BGC domains. This comparison showed that UProC was on average 18 times faster than HMMER (Supplementary Fig. 1a). We then evaluated the accuracy of BiG-MEx Operational Domain Unit (ODU) diversity estimation approach. We used BiG-MEx to compute the ODU diversity of the NRPS adenylation (AMP-binding) and condensation domains, as well as the PKS ketosynthase (PKS_KS) and acyltransferase (PKS_AT) domains in a simulated metagenomic dataset (Marine-TM dataset; see Materials and Methods section 3). Additionally, we computed the ODU diversity of these domains based on the domain sequences obtained from the genome sequences used to simulate the Marine-TM metagenomes. The latter estimates (henceforth, the reference estimates) were assumed to accurately reflect the ODU diversity, as they were computed using the complete domain sequences. We compared BiG-MEx ODU diversity estimates against the reference ODU diversity and observed that these were highly correlated: PKS_KS domains had a Pearson’s r of 0.77, while for the other domains the Pearson’s r was greater than 0.9 (Supplementary Fig. 1b). Lastly, we evaluated BiG-MEx’s BGC class abundance prediction module. We point out that although we modelled the abundance of a few BGC subclasses, we refer to all as BGC classes. For this analysis, we used two different simulated metagenomic datasets, one for training and the other for testing the BGC class abundance models (Marine-RM and Marine-TM, respectively) (see Supplementary Table 1). We predicted the BGC class abundances in the Marine-TM metagenomes, using BiG-MEx BGC class abundance prediction module, and additionally, computed the BGC class abundances based on the complete genome sequences used to simulate the Marine-TM metagenomes. Similarly as indicated previously, the latter abundances were taken as a reference to evaluate the accuracy of the predictions. We observed that the predicted vs. reference abundance comparison for 20 of the 23 BGC classes we modelled (i.e., the total number of classes detected in the Marine-RM training dataset) had a Pearson’s r correlation coefficient greater than 0.5 and a median unsigned error (MUE) lower than 0.25 (Supplementary Fig. 2). Figure 2a displays the scatter plots of this comparison for the NRPS, terpene, and type I and II PKS BGC classes. To benchmark BiG-MEx BGC class abundance prediction module, we compared its abundance predictions against the abundance estimates derived from running antiSMASH on assemblies of the Marine-TM metagenomes (hereafter referred to as the “assembly approach”). The plots in Figure 2b display the Pearson correlation coefficients and the unsigned error distributions with respect to the reference abundances comparing both approaches for the same four BGC classes mentioned above. All BGC class abundance models included in this analysis were considerably more accurate than the assembly approach (Supplementary Fig. 3).

**Fig. 2.**
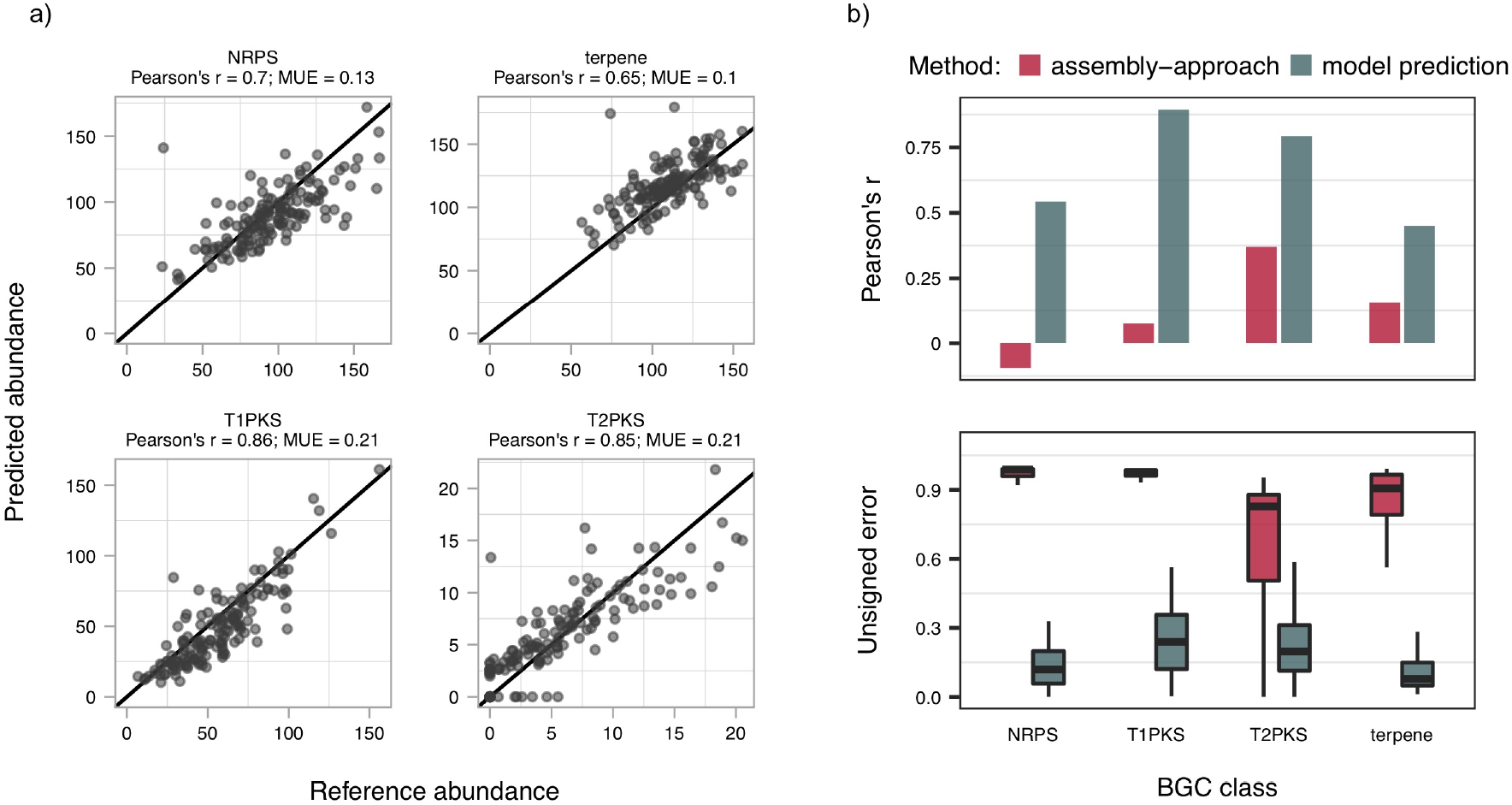
Evaluating and benchmarking the BGC abundance prediction models. (**a**) Scatter plots comparing the reference and predicted abundances of the NRPS, terpene, T1PKS and T2PKS BGC classes. MUE: Median Unsigned Error. The black, solid line represents the one-to-one relationship between the reference and predicted BGC class abundances. The BGC class abundance models were trained with the Marine-RM metagenomes and used to predict the abundances in the Marine-TM metagenomes. (**b**) Plots of the Pearson correlation coefficients (upper panel) and the unsigned error distributions (lower panel) of the BGC class abundances predicted by the models and estimated by the assembly approach, with respect to the reference abundances. In this comparison, we used 50 Marine-TM metagenomes. For the sake of clarity, 12 outlying unsigned error values (3% of the total comparisons) were excluded from the plot. The assembly approach consisted of the following tasks: 1) Assembling the metagenomes of the Marine-TM dataset; 2) Selecting the contigs with potential BGC sequences using BiG-MEx domain identification module; 3) Annotating the contigs with antiSMASH; 4) Mapping the short-read sequences to the identified BGC sequences; 5) Estimating the BGC class abundances.

To illustrate the application of BiG-MEx, we performed a Principal Coordinates Analysis (PCoA) based on BiG-MEx-derived BGC class abundance profiles of the 139 prokaryotic metagenomes of TARA Oceans. In Figure 3a, we ordinate the first two axes of the PCoA. The first axis (PCo1; 73.5% of the total variance) differentiated the mesopelagic (MES) from the surface (SRF) and deep chlorophyll maximum (DCM) water layers (Wilcoxon rank sum test; all p-values < 0.0001; see Supplementary Table 2). Further, the ordination values of the metagenomes along the PCo1 axis correlated with temperature (Pearson’s r = − 0.73; p-value < 0.0001). The differences in the BGC class composition between water layers were additionally confirmed with a Permutational Multivariate Analysis of Variance (PERMANOVA) (see Supplementary Table 3). We also performed a PCoA to explore the BGC domain composition and obtained a similar ordination of the metagenomes (Supplementary Fig. 4). These results are in agreement with previous work showing the stratification of microbial communities along depth and temperature gradients^19,20^. In particular, a very similar differentiation of the MES water layer along the first axis was also observed in the PCoA performed by Sunagawa et al.,^19^ based on the 16S _mi_tag (i.e., 16S ribosomal RNA gene tags^21^) composition of these same TARA Ocean metagenomes.

**Fig. 3.**
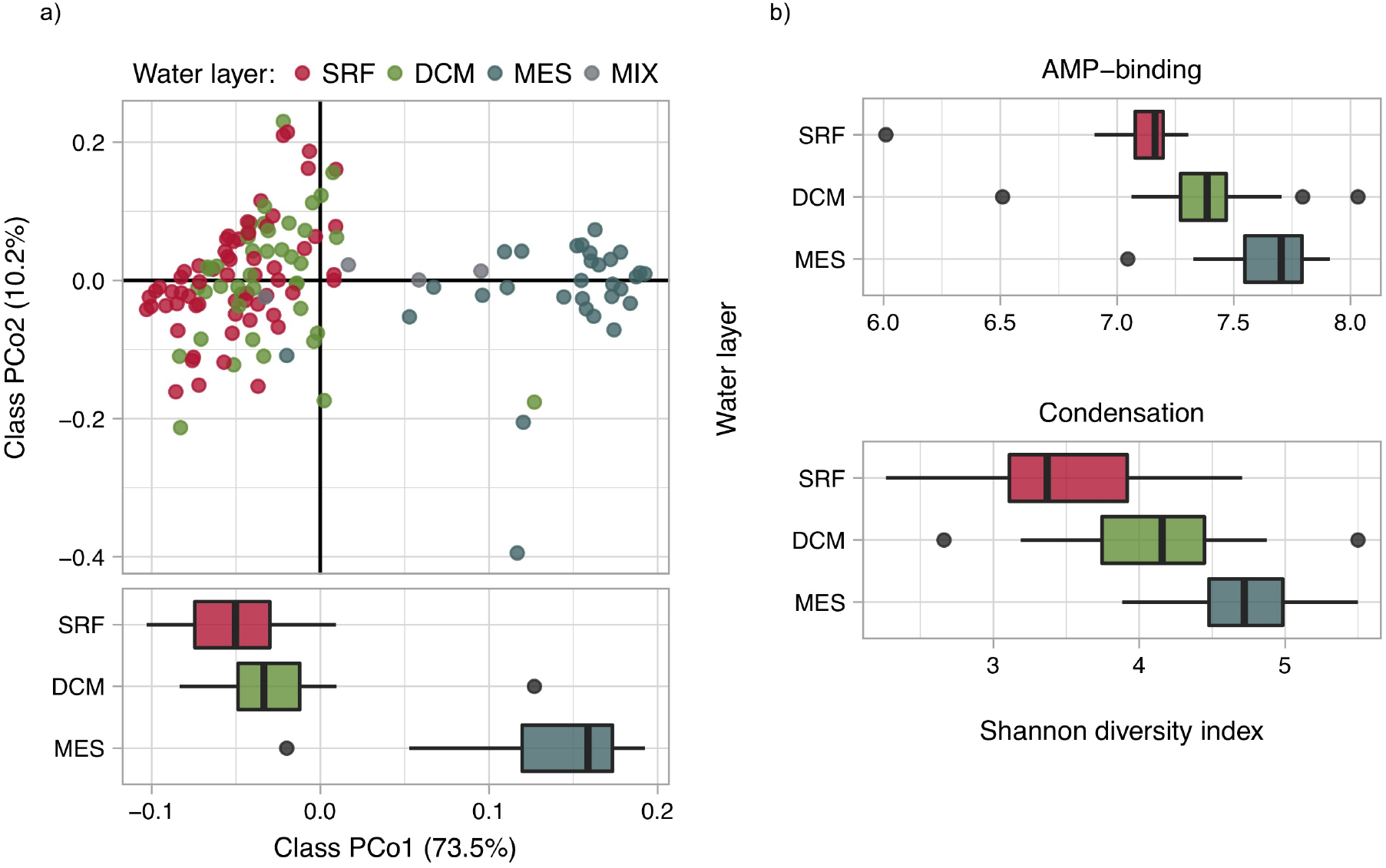
BiG-MEx BGC class composition and domain-based diversity analysis in the TARA Oceans dataset. **(a)** Principal Coordinates Analysis (PCoA) performed on a Bray-Curtis dissimilarity matrix of BGC class relative abundance profiles of the 139 prokaryotic metagenomes of TARA Oceans. BGC class abundance profiles were generated with BiG-MEx BGC class abundance module, using machine-learning models trained with the simulated Marine-RM metagenomic dataset. The abbreviations SRF, DCM, MES, and MIX correspond to surface, deep chlorophyll maximum, mesopelagic, and mixed epipelagic water layers, respectively. The boxplot in the bottom section of the panel shows the PCo1 value distributions for the metagenomes from the SRF, DCM and MES water layers. The PCo1 axis differentiated the MES water layer from the other two layers (Wilcoxon rank sum test; all p-values < 0.0001). **(b)** Bar plots showing the distribution of the ODU Shannon alpha diversity indices for the AMP-binding and condensation domains (NRPSs). The ODU diversity was computed for a match subset of 63 TARA Oceans metagenomes representing SRF, DCM, and MES water layers in 21 sampling stations. The AMP-binding and Condensation ODU diversity estimates were significantly different between the three water layers (pairwise Wilcoxon rank sum test; all p-values < 0.0001).

Next, we used BiG-MEx domain-based diversity module to compare the Operational Domain Unit (ODU) diversity of the NRPS adenylation (AMP-binding) and condensation domains between the SRF, DCM and MES water layers. These domains provide information about the chemical characteristics of the peptides synthesized by NRPS enzymes. AMP-binding domains recruit the amino acid monomers to be incorporated, while condensation domains catalyse the peptide bond formation^22,23^. In this analysis, we aimed to assess the potential chemical diversity of the NRPS products. NRPSs are one of the most studied BGC classes and are responsible for the production of many compounds with clinical applications. The results show that the ODU diversity of both domains increased from the surface to the mesopelagic water layers and differentiated significantly between water layers (pairwise Wilcoxon rank sum test; all p-values < 0.005; see Supplementary Table 2) (Fig. 3b). These results indicate that the microbial communities inhabiting deeper water layers contain a significantly higher diversity of NRPS products. The ODU diversity gradients resemble the Operational Taxonomic Unit (OTU) richness and functional diversity distributions shown in Sunagawa et al. We found highly significant correlations between the ODU diversity estimates and the taxonomic and functional richness and diversity obtained by Sunagawa et al. (see Supplementary Table 4).

To exemplify a more fine-grained analysis with BiG-MEx’s domain-based diversity module, we explored the ODU diversity of condensation domains in the three TARA Oceans metagenomes obtained from the SRF, DCM, and MES water layers at the sampling station TARA_085 (Antarctic Ocean). As observed previously, the metagenome from the MES water layer had a higher ODU diversity (Fig. 4a). It contains many low abundance ODUs scattered throughout the reference phylogeny (Fig. 4b). The phylogenetic diversity^24^ (PD) of ODU representative sequences of the MES metagenome, was 5.24 and 2.65 times greater than the PD estimates of the SRF and DCM metagenomes, respectively. Besides indicating a higher chemical diversity, this result indicates that there is greater potential chemical novelty of nonribosomal peptides. Additionally, the phylogenetic placement analysis revealed that the most abundant condensation ODU is placed close to the reference condensation domain sequences of NRPSs that produce albicidin and cystobactamide antibiotics (both topoisomerase inhibitors) (Fig. 4c). As albicidin is also a phytotoxin, the dominance of such ODU, which originates from the DCM layer, could be explained by the presence of a large number of NRPSs that act on the photosynthetic organisms that concentrate therein. The DCM layer had a notably higher chlorophyll concentration than the other two layers (0.01, 0.28, and 0 mg/m3 for the SRF, DCM, and MES respectively). The NRPS producing albicidin belongs to the class *Gammaproteobacteria* and order *Xanthomonadales*. This is in agreement with the ODU taxonomic affiliation, which was annotated as a *Gammaproteobacteria* (lowest common ancestor). This finding is also supported by the fact that the BLASTP search against the reference MIBiG database, showed that condensation domains significantly similar to NRPS domains producing albicidin (e-value < 1e-5), where only found in the DCM layer. We cannot exclude other possible explanations of these results; however, this line of exploration might be worth considering for further research. Rising ocean temperatures, as a consequence of global warming, are predicted to increase the frequency of events of bacteria affecting the algae populations, which in turn can impact marine ecosystems on a global scale^25^. Regarding potential biotechnological applications, these results are relevant for bioprospecting, given that albicidin and cystobactamide are antibiotics of interest for clinical treatments^26,27^.

**Fig. 4.**
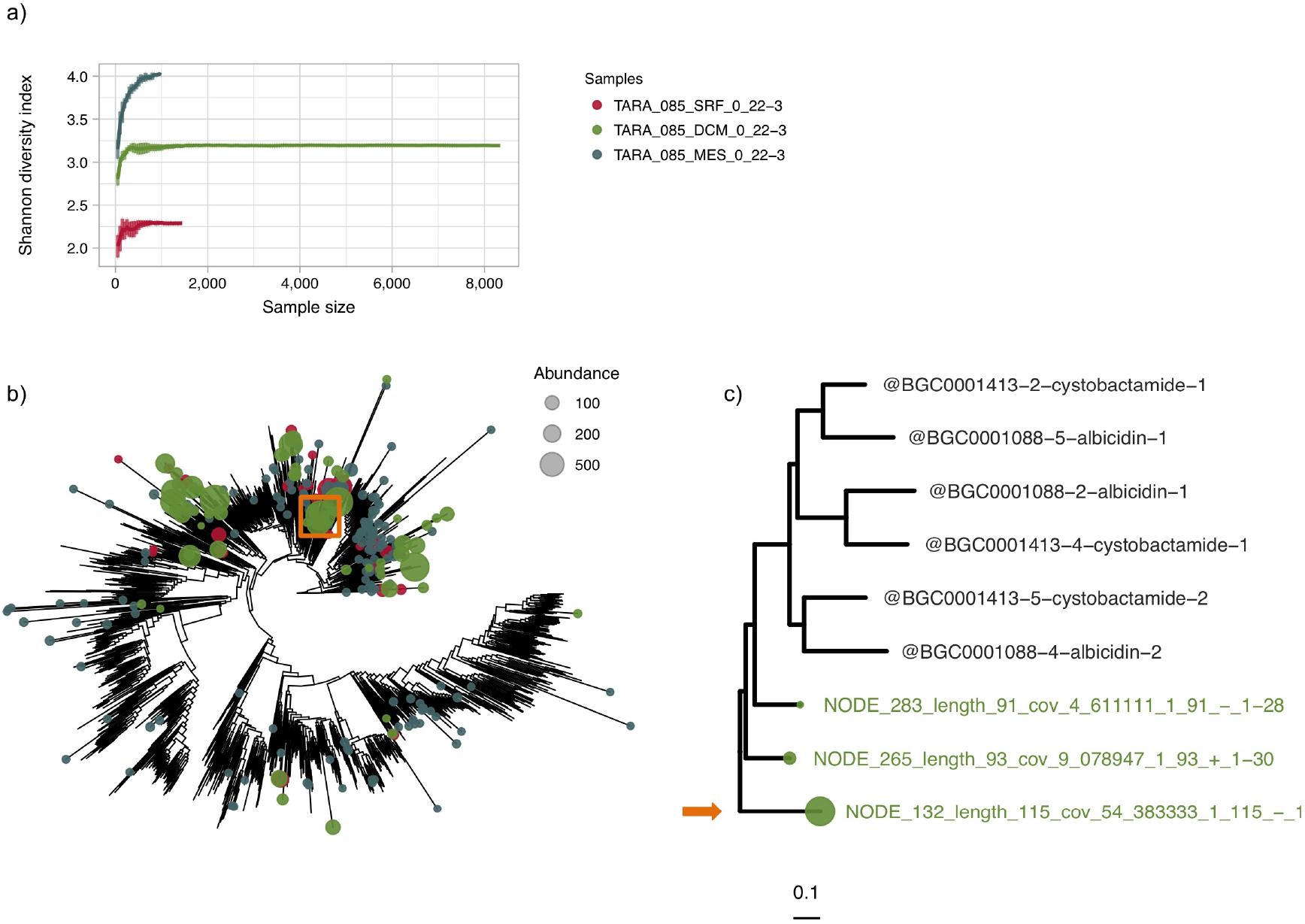
BiG-MEx diversity analysis of condensation domains in three metagenomes from TARA Oceans sampling station TARA_085. **(a)** Rarefaction curves of the Shannon alpha diversity indices generated by BiG-MEx domain-based diversity analysis module, comparing the diversity of condensation ODUs in the metagenomes of the SRF, DCM, and MES water layers. Condensation domain sequences were clustered into ODUs using a 75% amino acid identity threshold. The diversity was computed using the number and abundance of distinct condensation ODUs. **(b)** Phylogenetic placement of the condensation ODU representative sequences, as performed by the BiG-MEx domain-based diversity analysis module. The SRF, DCM and MES had a phylogenetic diversity (Faith’s PD)^24^ of 58.15, 114.98 and 304.88, respectively. The size and colour of the bubbles on the leaves represent the ODU abundance and sample origin, respectively. **(c)** Detail of the clade contained in the orange, hollow square in (c), including the most abundant ODU (obtained in the TARA_085_DCM_0_22-3 sample; indicated with an orange arrow).

We note that neither the TARA Oceans Metagenomes Assembled Genomes (MAGs)^28^, nor the DCM assembled metagenome from TARA_085 sampling site, contained albicidin or cystobactamide NRPS-like sequences. The difference between our findings in comparison to standard approaches based on assembled data was expected to occur, given the limitations of the latter to identify BGC classes (as shown in Fig. 2). In Supplementary Figure 5, we illustrate this problem by comparing the sequence length between MIBiG BGCs, and the TARA Oceans MAGs, and assembled metagenomic contigs.

Considering the relevance of human microbiome-derived natural product BGCs in medical research, we demonstrate the applicability of BiG-MEx to explore the BGC composition in the Human Microbiome Project (HMP) dataset. Our analyses traversed metagenomes from the buccal mucosa, tongue dorsum, and supragingival plaque body sites as well as stool samples (491 metagenomes in total). We used BiG-MEx to compute the BGC domain and class abundance profiles, and applied the same methodology as described for TARA Oceans, to compute the domain and class-based PCoAs. In agreement with previous analyses based on the taxonomic and functional annotation^4,29^, we observed that metagenomes grouped according to the body site they were sampled from in the first two ordination axes (Supplementary Fig. 6a and b). We conducted a PERMANOVA to test and assess the strength of the differences between body sites according to their BGC class composition, which showed significant differences in all body site comparisons (Supplementary Table 5). Additionally, we used BiG-MEx to compare the ODU diversity of the AMP-binding and condensation domains between body sites and observed that supragingival plaque metagenomes contain significantly higher diversity than the other body sites (pairwise Wilcoxon rank sum test; p-value < 0.0001) (Supplementary Figure 7 and Supplementary Table 6). This is in line with previous work showing that the supragingival plaque is one of the most functionally and taxonomically diverse body sites in the HMP dataset^4^.

Besides the mining analyses, BiG-MEx BGC class profiling can be used for the screening and prioritization of (meta)genomic samples. BGC class abundance profiles derived from shallow sequencing depth (meta)genomic data can be used for the identification of strains or environments with high biosynthetic potential, before investing in deep sequencing or long read sequencing technologies. As a proof-of-concept for this application, in Supplementary Fig. 8 we show a comparison of the BGC class abundance predictions computed in metagenomes of 100 and 5 million reads.

In our example applications, we processed 630 metagenomes, which sum to more than 85 billion paired-end reads. The analyses showed that BiG-MEx ultrafast domain and class profiling, and ODU diversity estimates provide biologically meaningful information, which can be used to mine BGCs in metagenomic data and as a basis from which to assess the ecological roles of their products in specific environments.

BiG-MEx extends BGC-based research and exploitation into large environmental datasets. It can be used to study the biogeography, distribution, and diversity of natural product BGCs either at the class, domain or ODU levels. Such analyses have the potential to accelerate the discovery of new bioactive products.

## Materials and Methods

### 1. Data acquisition, pre-processing and annotation

We retrieved the 139 prokaryotic metagenomes of the TARA Oceans dataset from the European Nucleotide Archive^30^ (ENA:PRJEB1787, filter size: 0.22-1.6 and 0.22-3). To pre-process the metagenomic short-read data, we clipped the adapter sequences (obtained from Shinichi Sunagawa personal communication, July 21, 2015) using the BBDuk tool from the BBMap 35.00 suite (https://sourceforge.net/projects/bbmap/) with a maximum Hamming distance of one (hdist=1). We then merged the paired-end reads using VSEARCH 2.3.4^31^, quality trimmed all reads at Q20 and filtered out sequences shorter than 45bp using BBDuk, and de-replicated the resulting quality-controlled sequences with VSEARCH. We annotated the BGC domains by first predicting the Open Reading Frames (ORFs) in the pre-processed data with FragGeneScan-plus^32^ and then running BiG-MEx on the predicted ORF’s amino acid sequences.

We downloaded 491 human microbiome metagenomes from the Data Analysis and Coordination Center (DACC) for the Human Microbiome Project (HMP) (https://www.hmpdacc.org/hmp/HMASM/). Our dataset included the metagenomes of the supragingival plaque (118), tongue dorsum (128), buccal mucosa (107), and the stool (138) body sites. These metagenomes have been already pre-processed as described in The Human Microbiome Project Consortium 2012^33^. The additional pre-processing tasks we performed consisted of merging the metagenomic reads with VSEARCH, quality trimming all reads at Q20 and filtering out sequences shorter than 45 bp with BBduk. To annotate the BGC domains, we predicted the ORFs with FragGeneScan-plus and ran BiG-MEx BGC domain identification module on the ORF’s amino acid sequences (Supplementary Table 7).

### 2. Exploratory analysis performed on TARA Oceans and HMP datasets

The domain abundance profiles of the TARA Oceans and HMP metagenomes were used to predict the BGC class abundance profiles with BiG-MEx BGC class abundance prediction module. The models used to generate the predictions for the TARA Oceans, and the oral and stool HMP metagenomes, were trained with the Marine-RM, Human-Oral and Human-Stool simulated metagenomic datasets, respectively. For each dataset, we performed a Principal Coordinate Analysis (PCoA) as follows: 1) We applied a total sum scaling standardization to both the domain and class abundance matrices; 2) We used the standardized matrices to compute the domain and class Bray-Curtis dissimilarity matrices; 3) We performed the PCoAs on the dissimilarity matrices with vegan R package utilizing the function capscale^34^.

We applied a Permutational Multivariate Analysis of Variance (PERMANOVA)^35^ to quantify the strength and test the differences between water layers and body sites according to their BGC class composition. For these analyses, we selected a balanced subset of metagenomes from the TARA Oceans and HMP datasets (63 and 216 metagenomes, respectively; see below). We performed a PERMANOVA on the Bray-Curtis dissimilarity matrix, computed for the TARA Oceans and HMP metagenome subsets as described above, to test the differentiation between all groups simultaneously. Subsequently, we tested each pair of groups independently, applying the Bonferroni correction for multiple comparisons. To perform the PERMANOVA, we employed the adonis function of the vegan R package, with the permutation parameter set to 999.

To compare the domain ODU diversity of the NRPS adenylation (AMP-binding) and condensation domains between the surface (SRF), deep chlorophyll maximum (CDM) and mesopelagic (MES) water layers, we used a subset of 63 TARA Oceans metagenomes, representing the three water layers in 21 sampling stations. We computed the ODU Shannon diversity in these metagenomes, using routines implemented in the BiG-MEx domain-based diversity module. Additionally, we used the same BiG-MEx module to examine the diversity of the condensation domains in the metagenomes representing the three water layers at sampling station TARA_085. To perform the ODU taxonomy annotation, we used MMseqs2 taxonomy assignment function^36^ based on UniRef100^37^ sequences (release-2018_08), with the e-value and sensitivity parameters set to 0.75 and 0.01, respectively. To compare the AMP-binding and condensation ODU diversity between body sites, we applied a similar approach as described above. We selected a subset of 216 metagenomes, 54 from each of the supragingival plaque, tongue dorsum, buccal mucosa, and stool body sites. This subset includes only the metagenomes obtained from individuals for whom the four body sites were sampled. We applied BiG-MEx domain-based diversity module to compute the ODU Shannon diversity estimates.

The Wilcoxon rank-sum tests (two-sided) to assess the significance of the differentiations between metagenomes from different groups (i.e., water layers or body sites), were performed with the wilcox.test function from the R package stats^38^.

### 3. Data simulation, pre-processing and annotation

#### 3.1 Construction of simulated metagenomic datasets

We created four simulated metagenomic datasets: Two of these approximate the taxonomic composition found in marine environments (Marine-RM and Marine-TM), and the other two, the taxonomic composition found in the human oral cavity and stool body sites (Human-Oral and Human-Stool, respectively). Each dataset is composed of 150 metagenomes, all of which have a size of two million paired-end reads. To simulate a metagenomic dataset, we first created a dataset of reference genome sequences and the genome abundance profiles to specify the metagenomes’ taxonomic composition. That is, we defined a hypothetical microbial community from which a metagenome is simulated by specifying which reference genomes and the number of times each genome occurs in the community.

To create the Marine-RM (Marine Reference Microbiome) genome dataset, we downloaded all genomes belonging to the Ocean Microbial Reference Gene Catalogue (OM-RGC)^19^ having an assembly status of “Complete genome” from RefSeq^39^ (on December 7^th^, 2017). If a given species did not have a complete genome sequence available, we randomly selected another species of the same genus. In total, we obtained 378 genomes corresponding to 363 species.

We applied a similar methodology to create the Marine-TM (Marine TARA Microbiome) genome dataset. To determine the taxonomic composition, we used the genus affiliation of TARA Oceans Operational Taxonomic Units (OTUs)^19^. We only included 30 shared genera (randomly selected) between TARA OTU and the Marine-TM genome dataset. This latter filtering was necessary to reduce the taxonomic overlap, given that we used the Marine-TM dataset to evaluate the performance of the BGC class abundance models trained with the Marine-RM dataset (see section 4.3). For the remaining genera for which there was at least one representative completely sequenced genome, we downloaded a maximum of three genomes per genus from RefSeq, irrespective of their species affiliation. This resulted in a database composed of 344 genomes from 308 species.

To create the genome datasets for the Human-Oral and Human-Stool metagenomic datasets, we used the genomes sequenced by the HMP derived from samples of the oral cavity and stool body sites. Given that few of these genomes were completely sequenced, we also included partially complete sequenced genomes. We downloaded all genomes with an assembly status of “Complete genome” or “Chromosome” or “Scaffold” generated by the HMP from the GenBank database^40^ (on March 15th, 2018). In the cases where a genome (sequenced by the HMP) had an assembly status lower than “Scaffold”, we downloaded another genome with the same species affiliation and an assembly status of “Complete genome” or “Chromosome”. The Human-Oral and Human-Stool reference genome datasets contain 209, and 479 genomes representing 140 and 338 species, respectively.

To create the community abundance profile of a metagenomic dataset, we randomly selected between 20 and 80 genomes from its genome reference dataset and defined the number of times each genome occurs by sampling from a lognormal distribution with mean 1 and standard deviation of 0.5. Lastly, we simulated the metagenomes with MetaSim v0.9.5^41^. MetaSim was set to generate paired-end reads with a length of 101bp, and a substitution rate increasing constantly along each read from 1×10^−4^ to 9.9×10^−2^. With this data, we aimed to simulate the short-read sequences generated by an Illumina HiSeq 2000 platform.

Dataset statistics are shown in Supplementary Table 1. The assembly accessions, organism names, taxids and NCBI FTP paths of the genome sequences used to create the genome databases are found in the Supplementary File 1. The workflow used to create the simulated metagenomic datasets can be found at https://github.com/pereiramemo/BiG-MEx/wiki/Data-simulation

### 3.2 Annotation of the simulated metagenomes

To estimate the reference BGC class abundances in a simulated metagenome, we annotated the BGC classes in its reference genome sequences with antiSMASH 3.0, mapped the paired-end reads to the identified BGC sequences with BWA-MEM 0.7.12^42^, and filtered out read alignments with a quality score lower than 10. Next, we removed read duplicates with Picard tools v1.133 (http://broadinstitute.github.io/picard), and computed the mean coverage with BEDtools v2.23^43^. The coverage estimates were assumed to accurately reflect the BGC class coverage-based abundances, as they were computed using complete BGC sequences, obtained from the genome sequences used to simulate the metagenomes. Additionally, we merged the paired-end reads of the simulated metagenomes with VSEARCH 2.3.4, predicted the ORFs with FragGeneScan-plus, and used BiG-MEx domain identification module to annotate the BGC domains in the ORF’s amino acid sequences. The workflow to annotate the synthetic metagenomes can be found at https://github.com/pereiramemo/BiG-MEx/wiki/Data-simulation#7-bgc-domain-annotation

### 4. Performance evaluation

#### 4.1 BGC domain identification module

We compared the running time (wall-clock) of UProC (i.e., uproc-prot) against a typical search using hmmsearch from the HMMER3 package^18^, for the identification of the 150 BGC domains included in BiG-MEx, in nine prokaryotic metagenomes of the TARA Oceans dataset (Supplementary Table 8). To run hmmsearch, we used the domain HMM profiles of antiSMASH. We annotated the nine metagenomes with both these tools in four independent rounds, each round using a different thread number (i.e., 4, 8, 16 and 32 threads). All parameters of uproc-prot and hmmsearch were set to default. The annotations were carried out on a workstation with Intel(R) Xeon(R) CPU E7-4820 v4 2.00GHz processors.

#### 4.2 BGC domain-based diversity analysis module

We evaluated BiG-MEx Operation Domain Unit (ODU) diversity estimation approach using NRPS adenylation (AMP-binding) and condensation, and PKS ketosynthase and acyltransferase domains (PKS_KS and PKS_AT, respectively). In this analysis, we used the BGC domain-based diversity analysis module to compute the ODU diversity in the Marine-TM dataset, and compared these estimates with the ODU diversity computed using the complete domain sequences. To obtain the latter ODU diversity, we applied the workflow implemented in BiG-MEx, with the exception that instead of assembling the domain sequences, we extracted these from the complete genome sequences used to simulate the Marine-TM metagenomes. We annotated the four domains in the complete genome sequences with hmmsearch using the antiSMASH HMM profiles.

#### 4.3 BGC class abundance predictions

We used the BGC class models trained with the Marine-RM metagenomic dataset to predict the BGC class abundances in the Marine-TM metagenomic dataset. We applied the methodology described in section 3.2 to compute the BGC class abundances in the Marine-TM metagenomes based on the complete genome sequences (i.e., reference abundance). To predict the BGC class abundances using machine-learning models, we annotated the Marine-TM metagenomes with the BiG-MEx domain identification module and used the domain abundance profiles as an input for the BiG-MEx BGC class abundance prediction module. The evaluation consisted of computing the Pearson correlation and median unsigned squared error (MUE) between the predicted and reference BGC class abundances. The MUE was computed as | *Â* − *A* |*/ A*, where *Â* and *A* are the predicted and reference abundance, respectively. To benchmark the machine-learning models, we compared the BGC class abundance predictions against the abundance estimates based on the assembly of 50 metagenomes of the Marine-TM dataset (assembly approach). The assembly approach consisted of assembling the metagenomes with MEGAHIT (default parameters), running BiG-MEx domain identification module to select the contigs with potential BGC sequences, annotating the selected contigs with antiSMASH 3.0, and estimating the BGC class abundance following the same approach as described in section 3.2 (Supplementary Table 9). We computed the unsigned error, and the Pearson correlation coefficient of BGC class abundance estimates obtained by the assembly approach and predicted by BiG-MEx, with respect to the reference BGC class abundances. The analysis performed to evaluate the accuracy of the models can be reproduced here: https://rawgit.com/pereiramemo/BiG-MEx/master/machine_leaRning/bgcpred_workflow.html

#### 4.4 Evaluation of the BGC class abundance predictions in shallow metagenomes

We selected 30 merged pre-processed TARA Oceans metagenomes and randomly subsampled these to generate two sets of metagenomes, one with 100 million and the other with 5 million reads, using the seqtk v1.0 tool (https://github.com/lh3/seqtk). We then annotated the BGC domains and predicted the BGC class abundances in this data using BiG-MEx (as described in sections 1 and 2), and compared the BGC class abundance predictions between the two sets of metagenomes.

### 5. BiG-MEx implementation

#### 5.1 BGC domain identification module

BiG-MEx BGC domain identification module uses the UProC 1.2.0^16^ software to classify short-read sequences using BGC domain references. To train UProC for this purpose, we manually curated all amino acid sequences matching 150 antiSMASH hidden Markov model profiles (HMMs)^10^. In this task, we removed sequences shorter than 25 amino acids and checked for the presence of overlaps between sequences of different HMM profiles. In addition, we categorized multi-domain proteins into multiple families. For the training process, we included a set of negative control profiles to assess the ratio of false positive hits. Namely, we used the t2fas, fabH, bt1fas, ft1fas profiles as negative controls for the PKS_KS, t2ks, t2ks2, t2clf, Chal_sti_synt_N, Chal_sti_synt_C, hglD and hglE profiles. Once we curated the amino acid sequence data, we applied the SEG(mentation) low complexity filter from the NCBI Blast+ 2.2 Suite^44^ and created the UProC database. This UProC database can be downloaded from https://github.com/pereiramemo/BiG-MEx. Based on the identified reads containing a BGC domain sequence, the module computes a count-based abundance profile of BGC domains.

#### 5.2 BGC domain-based diversity analysis module

This module performs two different analyses: Operational Domain Unit (ODU) diversity estimation and phylogenetic placement of domain sequences. The pipeline to estimate the ODU diversity, analyses each domain independently, and consists of the following steps: 1) Short-read sequences, where the domain being studied was identified, are recruited to perform a targeted assembly metaSPAdes 3.11^45^ with default parameters; 2) The Open Reading Frames (ORFs) in the resulting contigs are predicted with FragGeneScan-Plus; 3) Domain sequences are identified within the ORF amino acid sequences with *hmmsearch* from HMMER v3 and extracted; 4) Domain amino acid sequences are clustered into ODUs using MMseqs2^46^ with the cascaded clustering option and the sensitivity parameter set to 7.5; 5) Annotated unassembled reads are mapped to the domain nucleotide sequences with BWA-MEM 0.7.12, and the mean depth coverage is calculated using BEDtools v2.23; 6) Based on this information, the coverage-based abundance of the ODUs is computed and used to estimate an ODU alpha Shannon diversity. To allow a comparison of the ODU diversity estimates between samples with different sequencing depth, we include an option to estimate the diversity for rarefied subsamples.

To perform the phylogenetic placement of domain sequences, we applied an approach similar to NaPDoS^11^. However, we extended the phylogenetic placement analysis to 48 domains and included more comprehensive reference trees, which are critical for the analysis of large metagenomic samples. In detail, the phylogenetic placement consists of aligning the target domain sequences to their corresponding reference multiple sequence alignment (MSA) with MAFFT^47^ (using --add option). Subsequently, the extended MSA together with its reference tree are used as the input to run pplacer^48^ (with parameters: --keep-at-most 10 and --discard-nonoverlapped; all other parameters set to default). pplacer performs the phylogenetic placement using the maximum-likelihood criteria and outputs the extended tree in Newick and jplace formats^49^, and a table with statistics and information about the placement of each sequence (i.e., likelihood, posterior probability, expected distance between placement locations (EDPL), pendant length, and edge number). To visualise the phylogenetic placement, a tree figure is generated using the ggtree R package^50^, where the coverage of the placed sequences is mapped on their tree tips and used to scale a bubble representation. Besides the phylogenetic placement, we included in this module an option to perform a BLASTP search of the assembled domain sequences against the reference domain sequences.

To construct the reference phylogenies, we first downloaded all the BGC amino acid sequences from the MIBiG database^51^. We identified the domain sequences with hmmsearch using the BGC domain HMM profiles from antiSMASH. Subsequently, we extracted and clustered these sequences with MMseqs2 to create a non-redundant dataset of amino acid sequences for each domain. If the number of reference sequences identified in the MIBiG database was greater than 500, we used a clustering threshold of 0.7 identity at the amino acid level; otherwise, the threshold was set to 0.9; all other parameters of MMseqs2 were set as specified previously. All domains with less than 20 representative sequences were discarded. This resulted in a subset of 48 domains that were considered for the phylogenetic reconstructions. For each set of domain representative sequences, we generated an MSA with MAFFT using the E-INS-I algorithm, removed sequence outliers with OD-seq^52^ and constructed a phylogenetic tree with RAxML^53^. To select the protein evolutionary model for the phylogenetic reconstruction, we used the automatic model selection implemented in RAxML with the maximum likelihood criterion. We used the GAMMA model of rate heterogeneity and searched the tree space using the rapid hill-climbing algorithm^54^, starting from a maximum parsimony tree. For the sake of reproducibility, we specified a random seed number (i.e., -p 12345). Finally, we used RAxML to root the trees and compute the SH-like support scores^55^. In Supplementary File 2, we provide for each domain phylogeny the number of sequences and amino acid substitution model used, the mean, standard deviation, maximum and minimum cophenetic distances between sequences, Faith’s phylogenetic diversity^24^ and the name of its corresponding BGC class.

#### 5.3 BGC class abundance prediction module

BiG-MEx uses machine-learning models to predict the abundance of the BGC classes, based on the counts of annotated domains in unassembled metagenomes. Each model is class-specific and was trained using the abundance of the BGC class and its corresponding protein domains, as the response and predictor variables, respectively. We used the classification rules defined in antiSMASH for the annotation of BGC classes, to determine the protein domains used as predictor variables in each model. To model the abundance of a given BGC class, we implemented a two-step zero-inflated process. First, the presence or absence of the target BGC class is predicted using a random forest (RF) binary classifier^56^. Second, a multiple linear regression (MLR) is applied to predict the class abundance, but only if the class was previously predicted as present. In the cases where the number of zero values was lower than 10 or non-existent, we directly applied an MLR. We trained the models using simulated metagenomic data (i.e., Marine-RM, Human-Oral and Human-Stool datasets). The models predict a coverage-based abundance since this was the response variable used in the training process. The RF binary classification models were created with the randomForest function of the randomForest R package^57^, with the parameters ntree set to 1000 (number of trees grown), nodesize set to 10 (minimum size of terminal nodes), and mtry set to 1 (number of variables randomly sampled as candidates at each split). For the MLR, we used the lm function of the stats R package (https://www.R-project.org/) with default parameters.

## Code availability

BiG-MEx is freely distributed using Docker container technology (www.docker.com), under the GNU General Public License v3.0. It can be downloaded from https://github.com/pereiramemo/BiG-MEx, where we also provide thorough documentation. Currently, we provide BGC class abundance models targeting the marine environment, four different human body sites, and the genus Streptomyces. To help users create their own BGC class abundance models and compute the predictions, we developed the R package bgcpred: https://github.com/pereiramemo/bgcpred. bgcpred is integrated in BiG-MEx, and is used to generate the BGC class abundance predictions.

## Data availability

In Supplementary file 1, we provide the GenBank and RefSeq assembly accessions for the genomes used to generate the simulated metagenomic datasets. We provide the BGC class and domain abundance tables, obtained from the simulated data, at https://github.com/pereiramemo/BiG-MEx/.

## Acknowledgements

EP-F was supported by a Ph.D. fellowship of the German Academic Exchange Service (DAAD) and the Uruguayan National Research and Innovation Agency (ANII). PLB was supported by the HGF Infrastructure Programme FRAM of the Alfred-Wegener-Institut, Helmholtz-Zentrum für Polar-und Meeresforschung. MHM was supported by Rubicon (825.13.001) and Veni (863.15.002) grants from the Netherlands Organization for Scientific Research (NWO). P-M was partially supported by German Research Foundation (DFG). AF-G received funding from the European Union’s Horizon 2020 research and innovation program [Blue Growth: Unlocking the potential of Seas and Oceans] under grant agreement no. [634486] 436 (project acronym INMARE). This work was accomplished using computational facilities provided by the Max Planck Society.

## Author contributions

EP-F, AF-G, PLB, and MHM conceived BiG-MEx’s algorithms. EP-F developed the tools, and analysed the data, and wrote the paper with contributions from all authors. All authors reviewed and approved the manuscript.

