## Supplementary figures for "Mining metagenomes for natural product biosynthetic gene clusters: unlocking new potential with ultrafast techniques"

### Slide 1
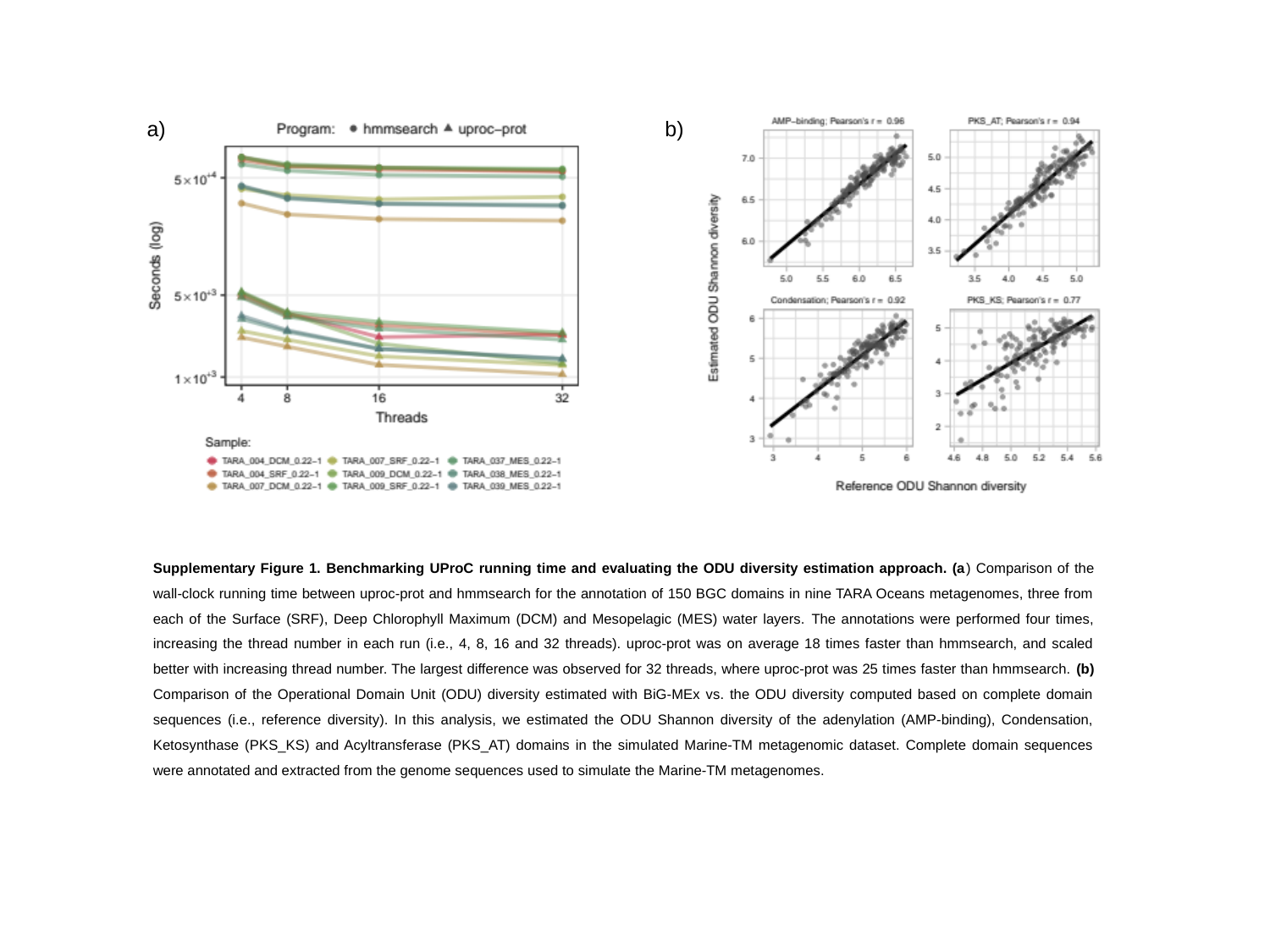

a)
b)
Supplementary Figure 1. Benchmarking UProC running time and evaluating the ODU diversity estimation approach. (a) Comparison of the wall‐clock running time between uproc-prot and hmmsearch for the annotation of 150 BGC domains in nine TARA Oceans metagenomes, three from each of the Surface (SRF), Deep Chlorophyll Maximum (DCM) and Mesopelagic (MES) water layers. The annotations were performed four times, increasing the thread number in each run (i.e., 4, 8, 16 and 32 threads). uproc-prot was on average 18 times faster than hmmsearch, and scaled better with increasing thread number. The largest difference was observed for 32 threads, where uproc-prot was 25 times faster than hmmsearch. (b) Comparison of the Operational Domain Unit (ODU) diversity estimated with BiG-MEx vs. the ODU diversity computed based on complete domain sequences (i.e., reference diversity). In this analysis, we estimated the ODU Shannon diversity of the adenylation (AMP-binding), Condensation, Ketosynthase (PKS_KS) and Acyltransferase (PKS_AT) domains in the simulated Marine-TM metagenomic dataset. Complete domain sequences were annotated and extracted from the genome sequences used to simulate the Marine-TM metagenomes.

### Slide 2
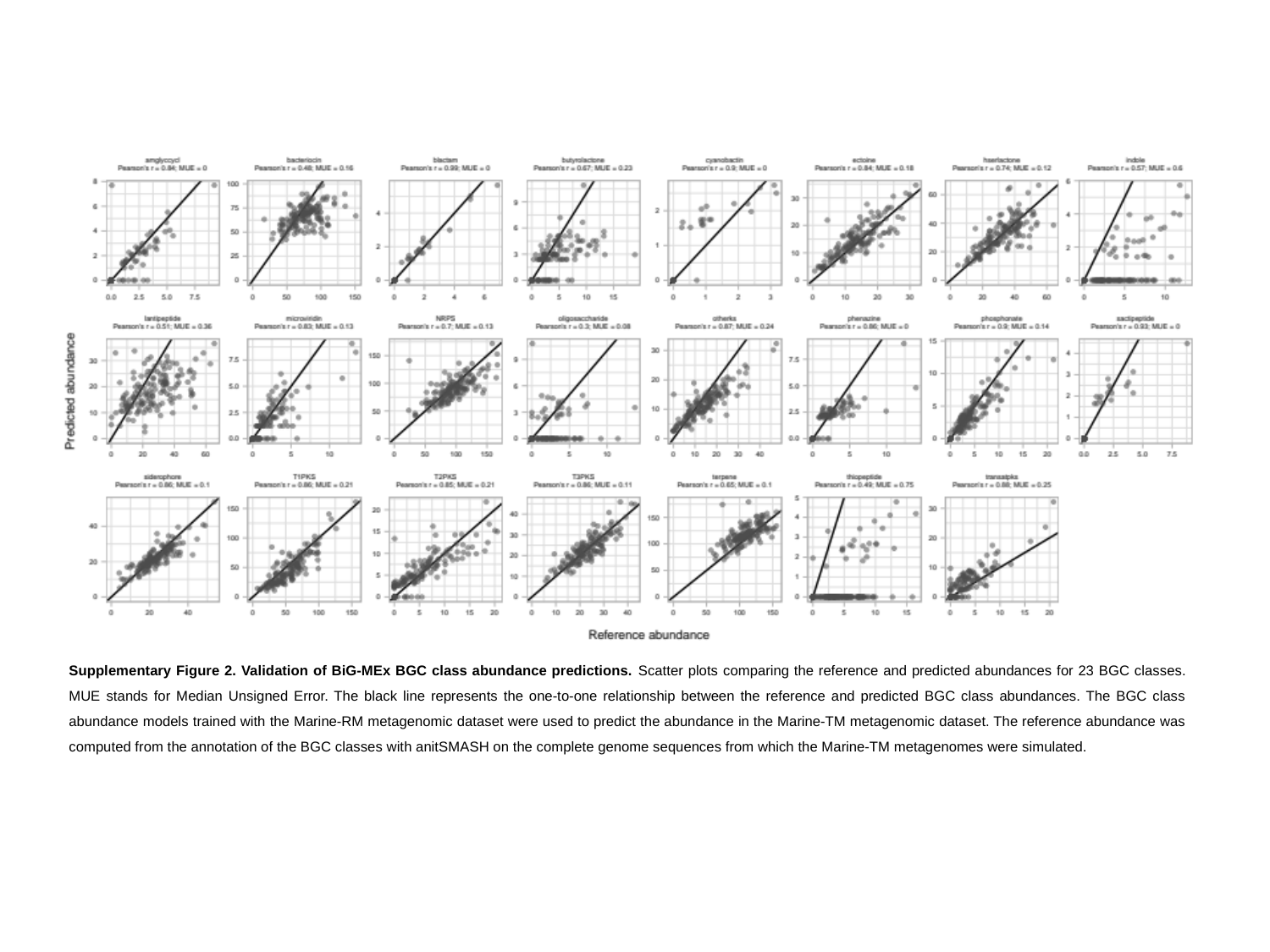

Supplementary Figure 2. Validation of BiG-MEx BGC class abundance predictions. Scatter plots comparing the reference and predicted abundances for 23 BGC classes. MUE stands for Median Unsigned Error. The black line represents the one-to-one relationship between the reference and predicted BGC class abundances. The BGC class abundance models trained with the Marine-RM metagenomic dataset were used to predict the abundance in the Marine-TM metagenomic dataset. The reference abundance was computed from the annotation of the BGC classes with anitSMASH on the complete genome sequences from which the Marine-TM metagenomes were simulated.

### Slide 3
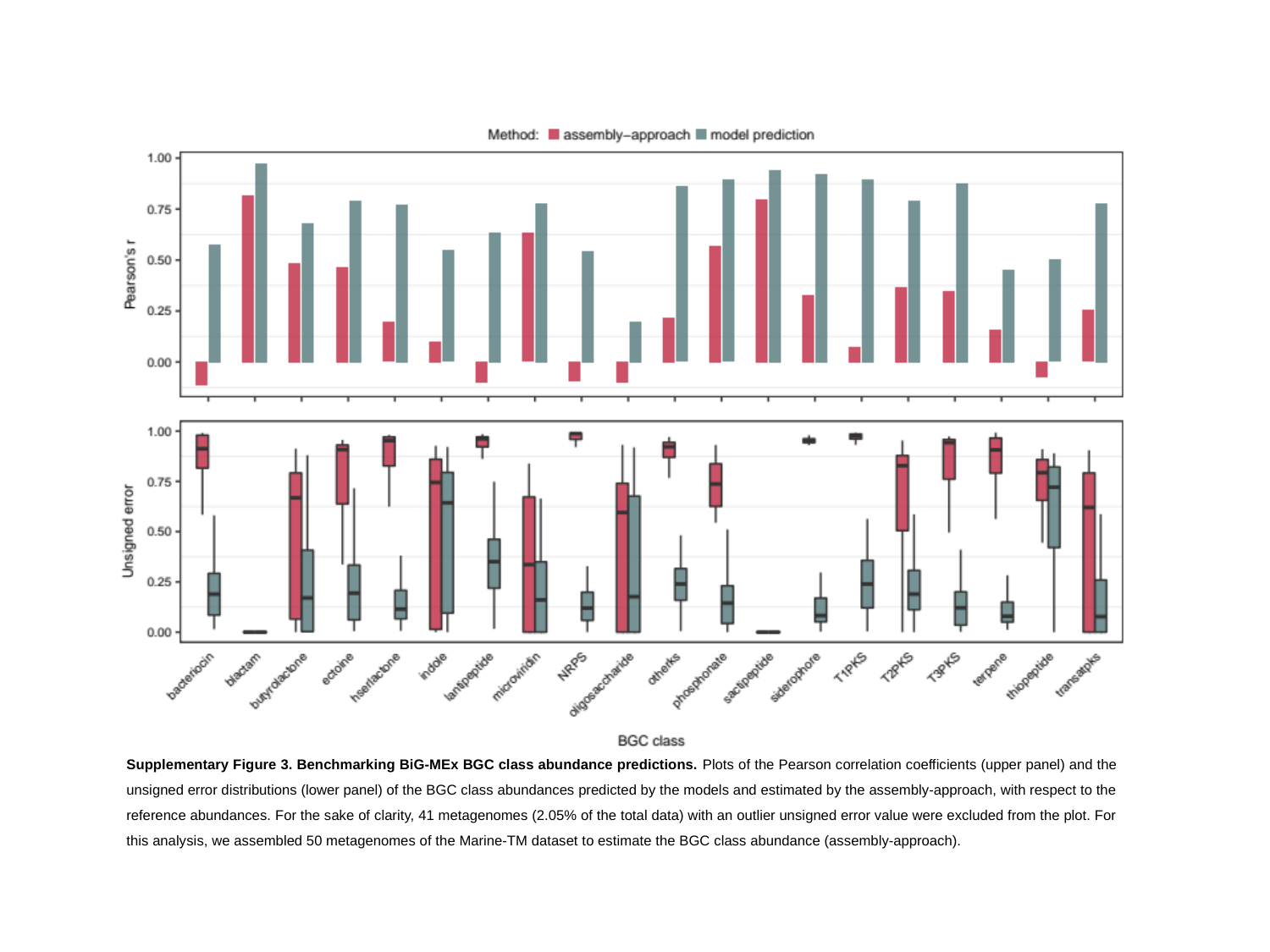

Supplementary Figure 3. Benchmarking BiG-MEx BGC class abundance predictions. Plots of the Pearson correlation coefficients (upper panel) and the unsigned error distributions (lower panel) of the BGC class abundances predicted by the models and estimated by the assembly-approach, with respect to the reference abundances. For the sake of clarity, 41 metagenomes (2.05% of the total data) with an outlier unsigned error value were excluded from the plot. For this analysis, we assembled 50 metagenomes of the Marine-TM dataset to estimate the BGC class abundance (assembly-approach).

### Slide 4
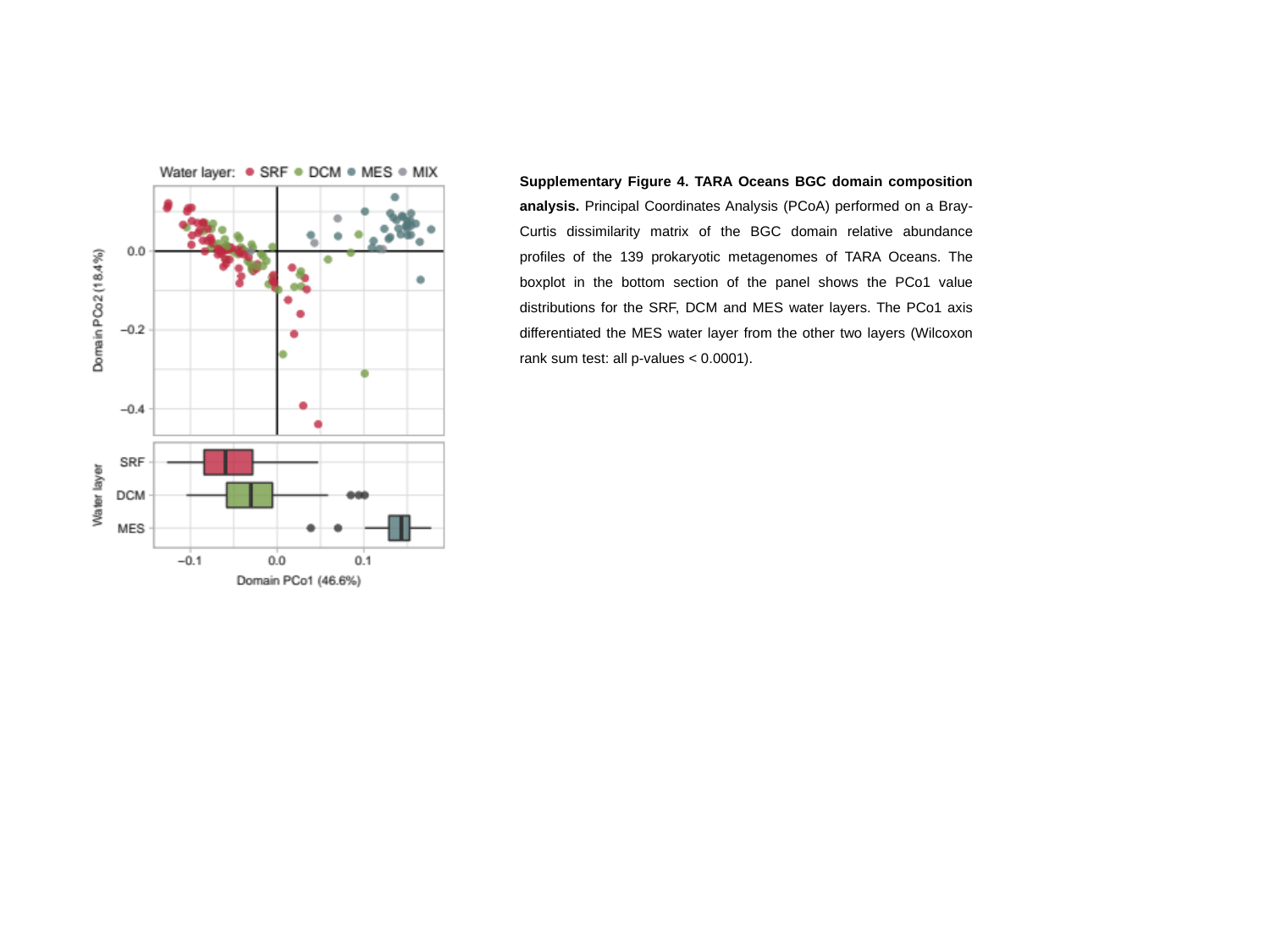

Supplementary Figure 4. TARA Oceans BGC domain composition analysis. Principal Coordinates Analysis (PCoA) performed on a Bray-Curtis dissimilarity matrix of the BGC domain relative abundance profiles of the 139 prokaryotic metagenomes of TARA Oceans. The boxplot in the bottom section of the panel shows the PCo1 value distributions for the SRF, DCM and MES water layers. The PCo1 axis differentiated the MES water layer from the other two layers (Wilcoxon rank sum test: all p-values < 0.0001).

### Slide 5
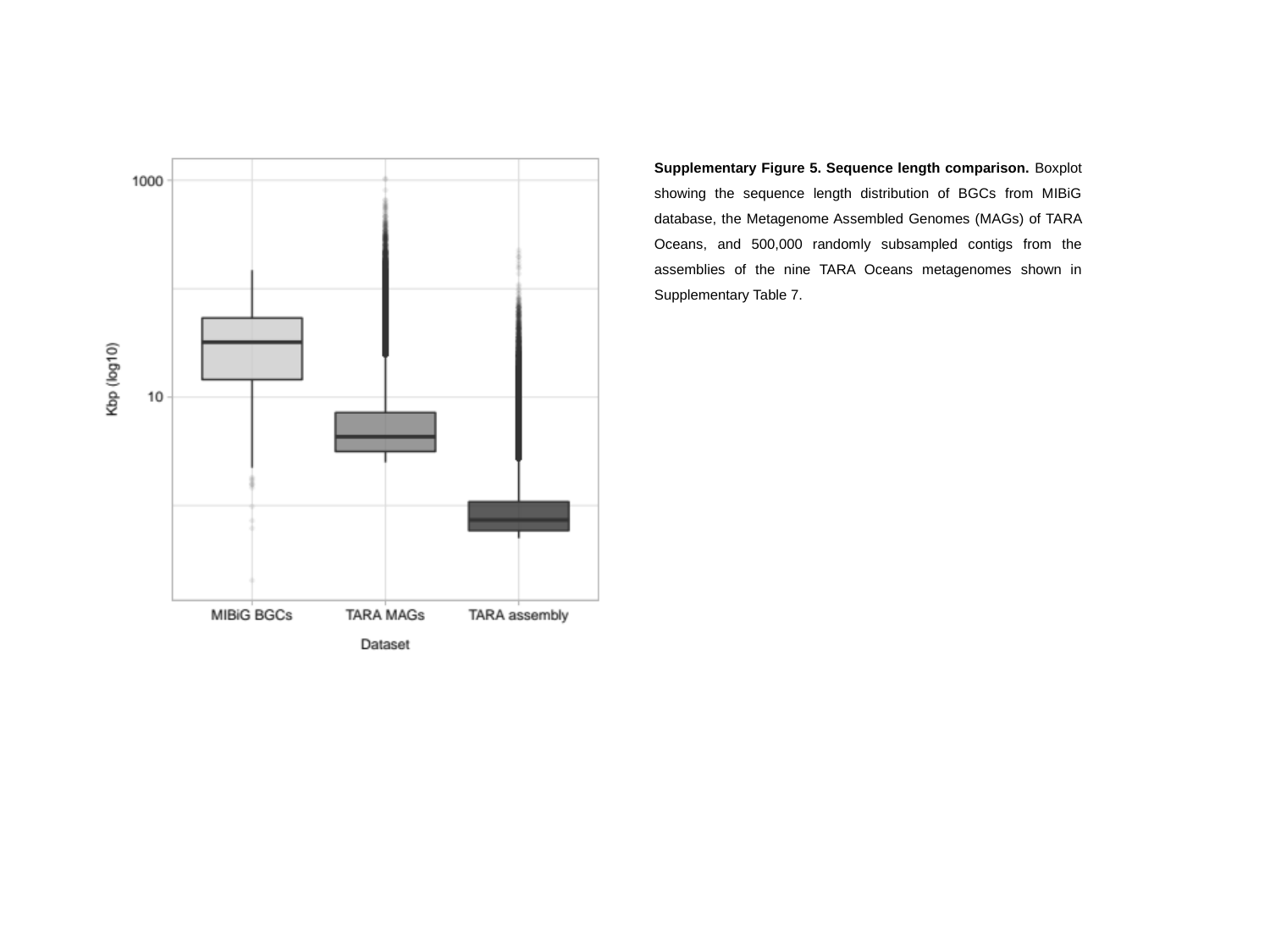

Supplementary Figure 5. Sequence length comparison. Boxplot showing the sequence length distribution of BGCs from MIBiG database, the Metagenome Assembled Genomes (MAGs) of TARA Oceans, and 500,000 randomly subsampled contigs from the assemblies of the nine TARA Oceans metagenomes shown in Supplementary Table 7.

### Slide 6
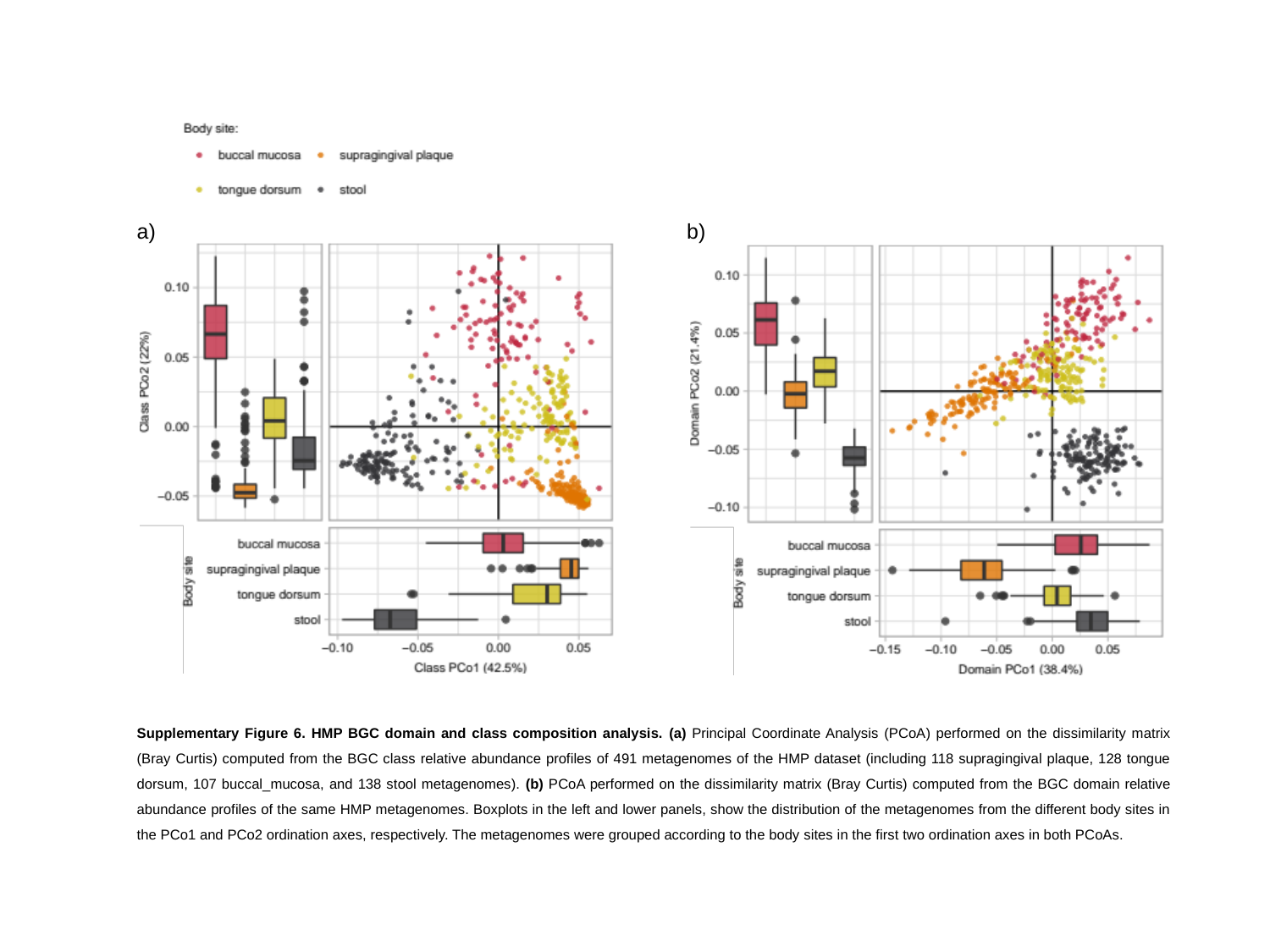

a)
b)
Supplementary Figure 6. HMP BGC domain and class composition analysis. (a) Principal Coordinate Analysis (PCoA) performed on the dissimilarity matrix (Bray Curtis) computed from the BGC class relative abundance profiles of 491 metagenomes of the HMP dataset (including 118 supragingival plaque, 128 tongue dorsum, 107 buccal_mucosa, and 138 stool metagenomes). (b) PCoA performed on the dissimilarity matrix (Bray Curtis) computed from the BGC domain relative abundance profiles of the same HMP metagenomes. Boxplots in the left and lower panels, show the distribution of the metagenomes from the different body sites in the PCo1 and PCo2 ordination axes, respectively. The metagenomes were grouped according to the body sites in the first two ordination axes in both PCoAs.

### Slide 7
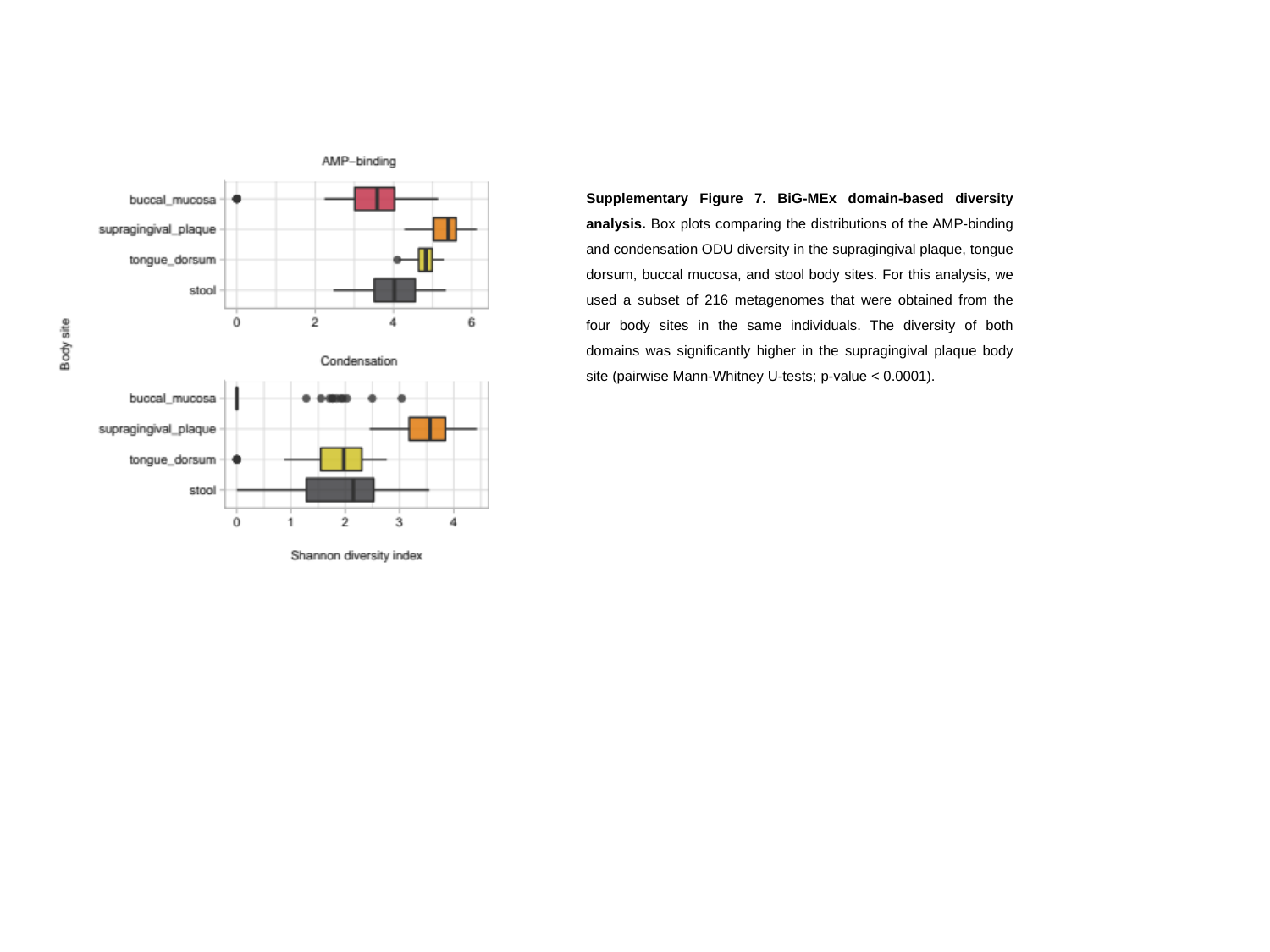

Supplementary Figure 7. BiG-MEx domain-based diversity analysis. Box plots comparing the distributions of the AMP-binding and condensation ODU diversity in the supragingival plaque, tongue dorsum, buccal mucosa, and stool body sites. For this analysis, we used a subset of 216 metagenomes that were obtained from the four body sites in the same individuals. The diversity of both domains was significantly higher in the supragingival plaque body site (pairwise Mann-Whitney U-tests; p-value < 0.0001).

### Slide 8
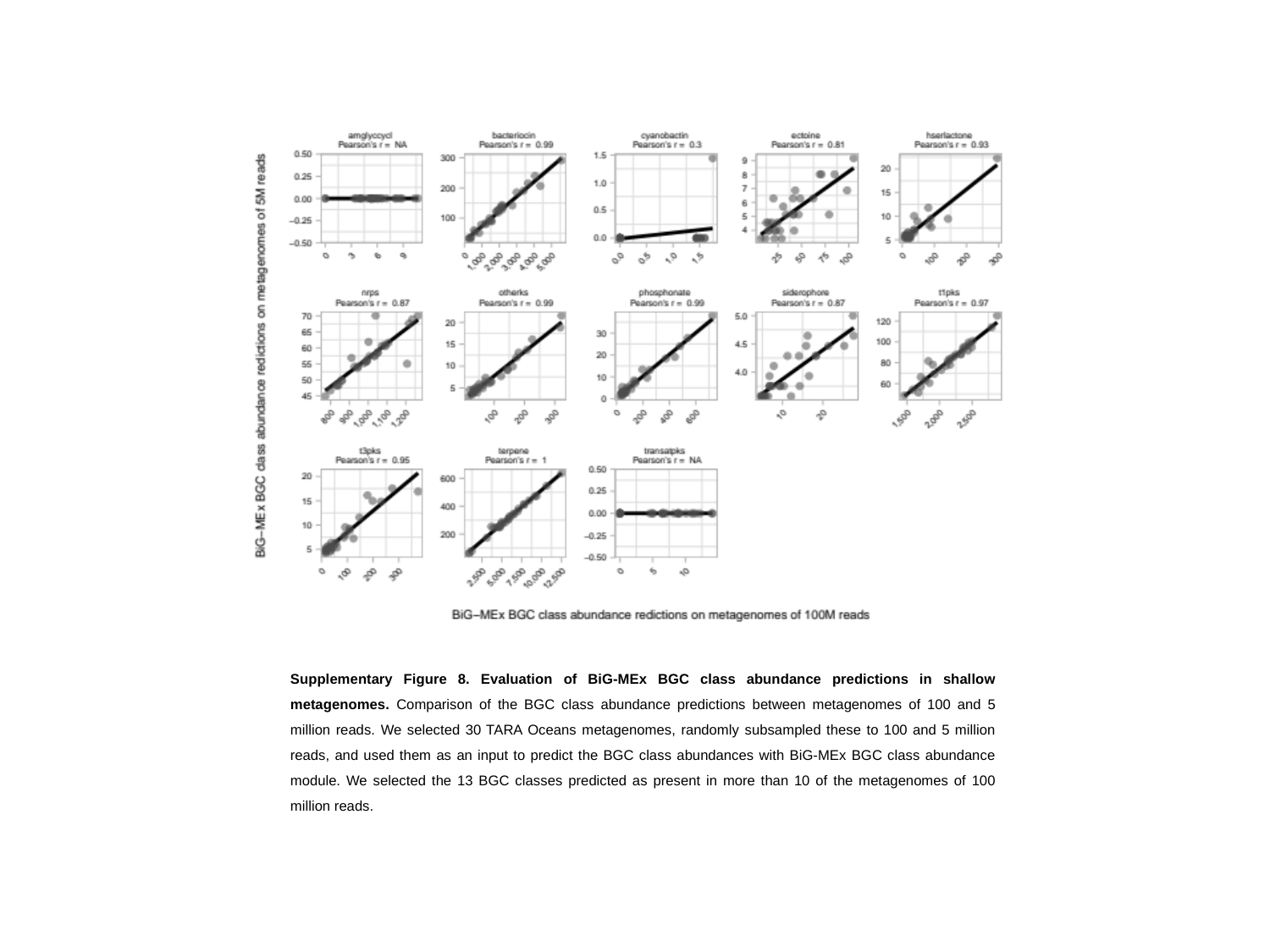

Supplementary Figure 8. Evaluation of BiG-MEx BGC class abundance predictions in shallow metagenomes. Comparison of the BGC class abundance predictions between metagenomes of 100 and 5 million reads. We selected 30 TARA Oceans metagenomes, randomly subsampled these to 100 and 5 million reads, and used them as an input to predict the BGC class abundances with BiG-MEx BGC class abundance module. We selected the 13 BGC classes predicted as present in more than 10 of the metagenomes of 100 million reads.
