## Supplementary tables for "Mining metagenomes for natural product biosynthetic gene clusters: unlocking new potential with ultrafast techniques"

### Slide 1
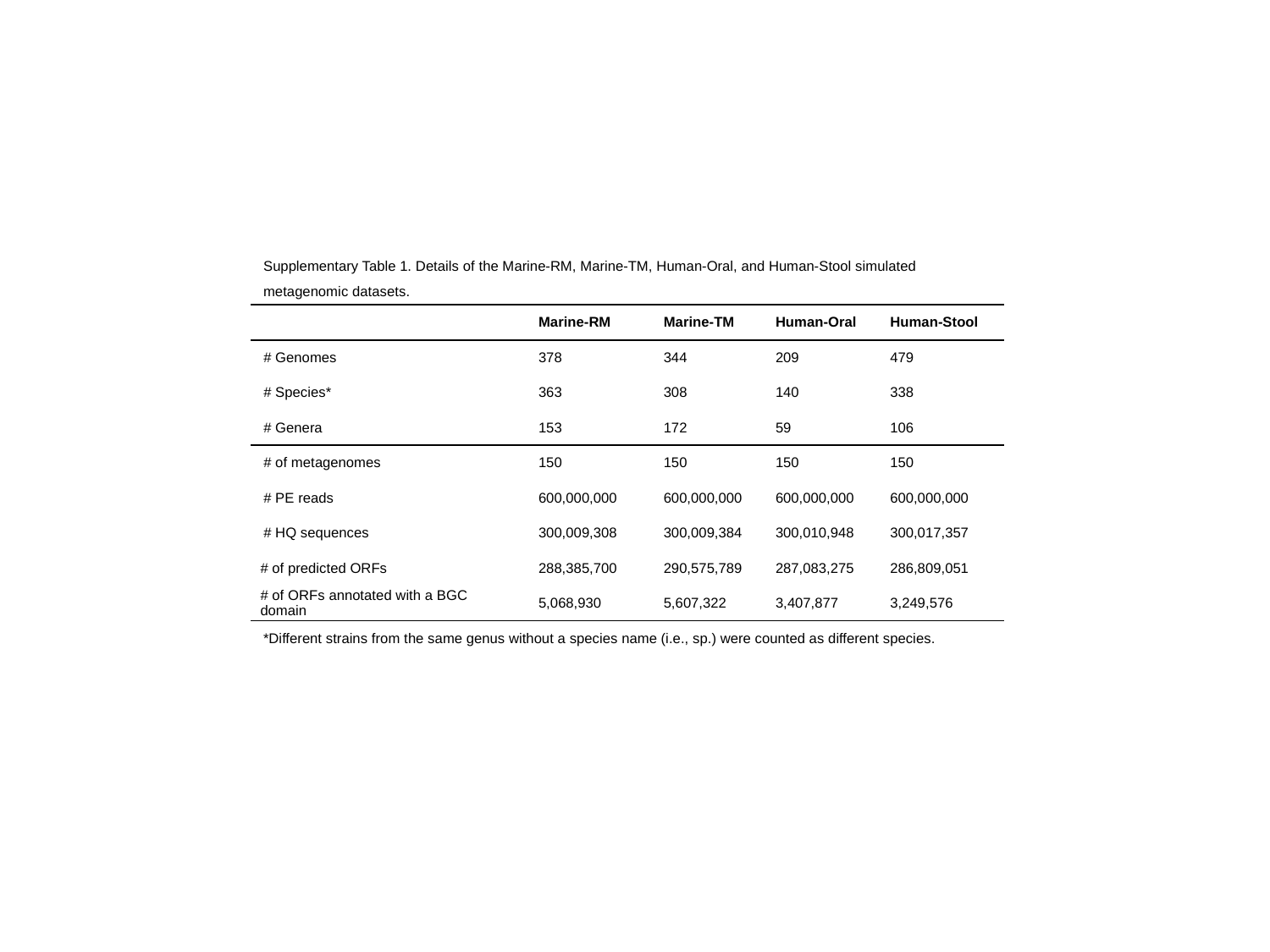

Supplementary Table 1. Details of the Marine-RM, Marine-TM, Human-Oral, and Human-Stool simulated metagenomic datasets.
| | Marine-RM | Marine-TM | Human-Oral | Human-Stool |
| --- | --- | --- | --- | --- |
| # Genomes | 378 | 344 | 209 | 479 |
| # Species\* | 363 | 308 | 140 | 338 |
| # Genera | 153 | 172 | 59 | 106 |
| # of metagenomes | 150 | 150 | 150 | 150 |
| # PE reads | 600,000,000 | 600,000,000 | 600,000,000 | 600,000,000 |
| # HQ sequences | 300,009,308 | 300,009,384 | 300,010,948 | 300,017,357 |
| # of predicted ORFs | 288,385,700 | 290,575,789 | 287,083,275 | 286,809,051 |
| # of ORFs annotated with a BGC domain | 5,068,930 | 5,607,322 | 3,407,877 | 3,249,576 |
*Different strains from the same genus without a species name (i.e., sp.) were counted as different species.

### Slide 2
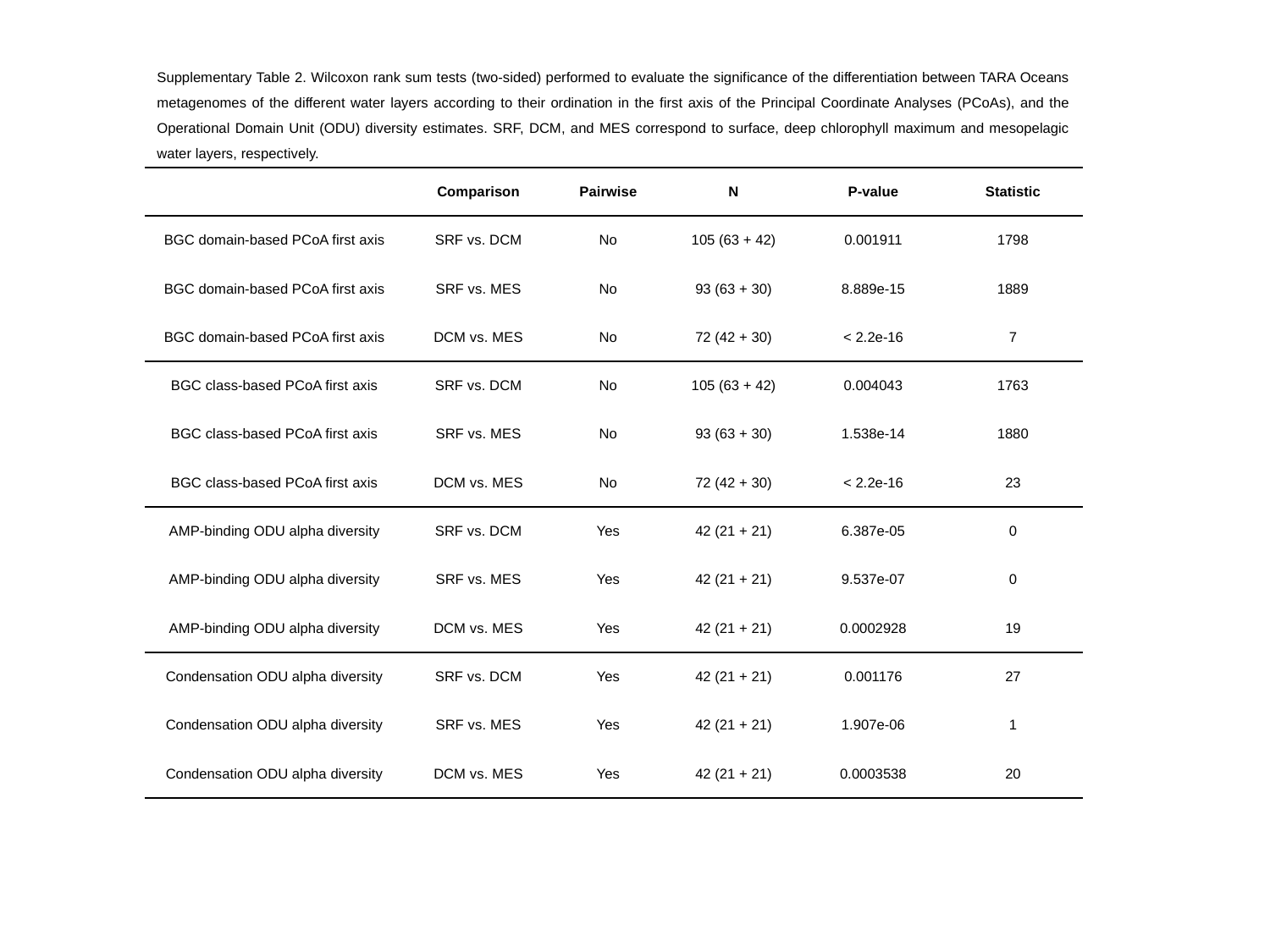

Supplementary Table 2. Wilcoxon rank sum tests (two-sided) performed to evaluate the significance of the differentiation between TARA Oceans metagenomes of the different water layers according to their ordination in the first axis of the Principal Coordinate Analyses (PCoAs), and the Operational Domain Unit (ODU) diversity estimates. SRF, DCM, and MES correspond to surface, deep chlorophyll maximum and mesopelagic water layers, respectively.
| | Comparison | Pairwise | N | P-value | Statistic |
| --- | --- | --- | --- | --- | --- |
| BGC domain-based PCoA first axis | SRF vs. DCM | No | 105 (63 + 42) | 0.001911 | 1798 |
| BGC domain-based PCoA first axis | SRF vs. MES | No | 93 (63 + 30) | 8.889e-15 | 1889 |
| BGC domain-based PCoA first axis | DCM vs. MES | No | 72 (42 + 30) | < 2.2e-16 | 7 |
| BGC class-based PCoA first axis | SRF vs. DCM | No | 105 (63 + 42) | 0.004043 | 1763 |
| BGC class-based PCoA first axis | SRF vs. MES | No | 93 (63 + 30) | 1.538e-14 | 1880 |
| BGC class-based PCoA first axis | DCM vs. MES | No | 72 (42 + 30) | < 2.2e-16 | 23 |
| AMP-binding ODU alpha diversity | SRF vs. DCM | Yes | 42 (21 + 21) | 6.387e-05 | 0 |
| AMP-binding ODU alpha diversity | SRF vs. MES | Yes | 42 (21 + 21) | 9.537e-07 | 0 |
| AMP-binding ODU alpha diversity | DCM vs. MES | Yes | 42 (21 + 21) | 0.0002928 | 19 |
| Condensation ODU alpha diversity | SRF vs. DCM | Yes | 42 (21 + 21) | 0.001176 | 27 |
| Condensation ODU alpha diversity | SRF vs. MES | Yes | 42 (21 + 21) | 1.907e-06 | 1 |
| Condensation ODU alpha diversity | DCM vs. MES | Yes | 42 (21 + 21) | 0.0003538 | 20 |

### Slide 3
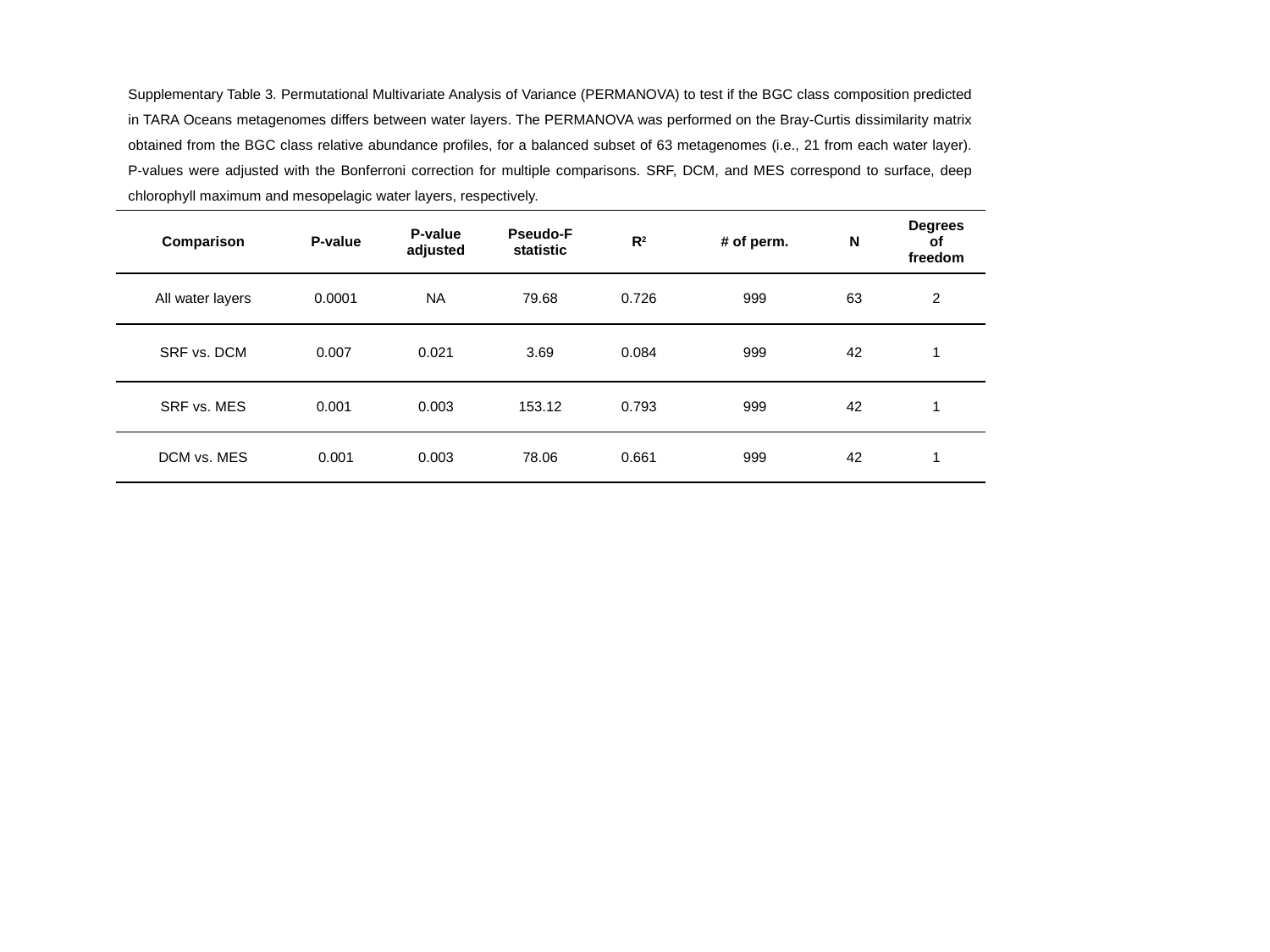

Supplementary Table 3. Permutational Multivariate Analysis of Variance (PERMANOVA) to test if the BGC class composition predicted in TARA Oceans metagenomes differs between water layers. The PERMANOVA was performed on the Bray-Curtis dissimilarity matrix obtained from the BGC class relative abundance profiles, for a balanced subset of 63 metagenomes (i.e., 21 from each water layer). P-values were adjusted with the Bonferroni correction for multiple comparisons. SRF, DCM, and MES correspond to surface, deep chlorophyll maximum and mesopelagic water layers, respectively.
| Comparison | P-value | P-value adjusted | Pseudo-F statistic | R^2^ | # of perm. | N | Degrees of freedom |
| --- | --- | --- | --- | --- | --- | --- | --- |
| All water layers | 0.0001 | NA | 79.68 | 0.726 | 999 | 63 | 2 |
| SRF vs. DCM | 0.007 | 0.021 | 3.69 | 0.084 | 999 | 42 | 1 |
| SRF vs. MES | 0.001 | 0.003 | 153.12 | 0.793 | 999 | 42 | 1 |
| DCM vs. MES | 0.001 | 0.003 | 78.06 | 0.661 | 999 | 42 | 1 |

### Slide 4
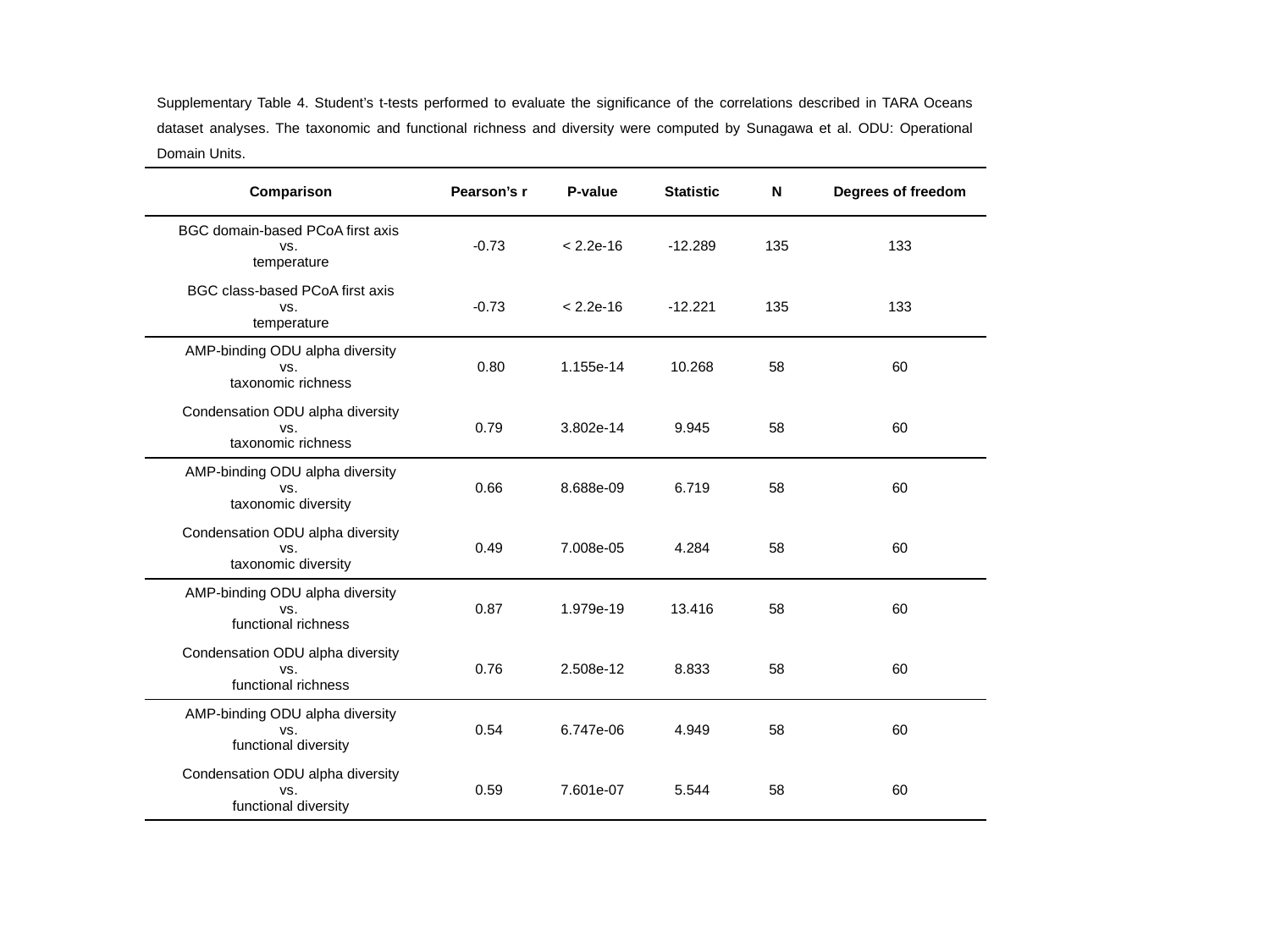

Supplementary Table 4. Student’s t-tests performed to evaluate the significance of the correlations described in TARA Oceans dataset analyses. The taxonomic and functional richness and diversity were computed by Sunagawa et al. ODU: Operational Domain Units.
| Comparison | Pearson’s r | P-value | Statistic | N | Degrees of freedom |
| --- | --- | --- | --- | --- | --- |
| BGC domain-based PCoA first axis vs. temperature | -0.73 | < 2.2e-16 | -12.289 | 135 | 133 |
| BGC class-based PCoA first axis vs. temperature | -0.73 | < 2.2e-16 | -12.221 | 135 | 133 |
| AMP-binding ODU alpha diversity vs. taxonomic richness | 0.80 | 1.155e-14 | 10.268 | 58 | 60 |
| Condensation ODU alpha diversity vs. taxonomic richness | 0.79 | 3.802e-14 | 9.945 | 58 | 60 |
| AMP-binding ODU alpha diversity vs. taxonomic diversity | 0.66 | 8.688e-09 | 6.719 | 58 | 60 |
| Condensation ODU alpha diversity vs. taxonomic diversity | 0.49 | 7.008e-05 | 4.284 | 58 | 60 |
| AMP-binding ODU alpha diversity vs. functional richness | 0.87 | 1.979e-19 | 13.416 | 58 | 60 |
| Condensation ODU alpha diversity vs. functional richness | 0.76 | 2.508e-12 | 8.833 | 58 | 60 |
| AMP-binding ODU alpha diversity vs. functional diversity | 0.54 | 6.747e-06 | 4.949 | 58 | 60 |
| Condensation ODU alpha diversity vs. functional diversity | 0.59 | 7.601e-07 | 5.544 | 58 | 60 |

### Slide 5
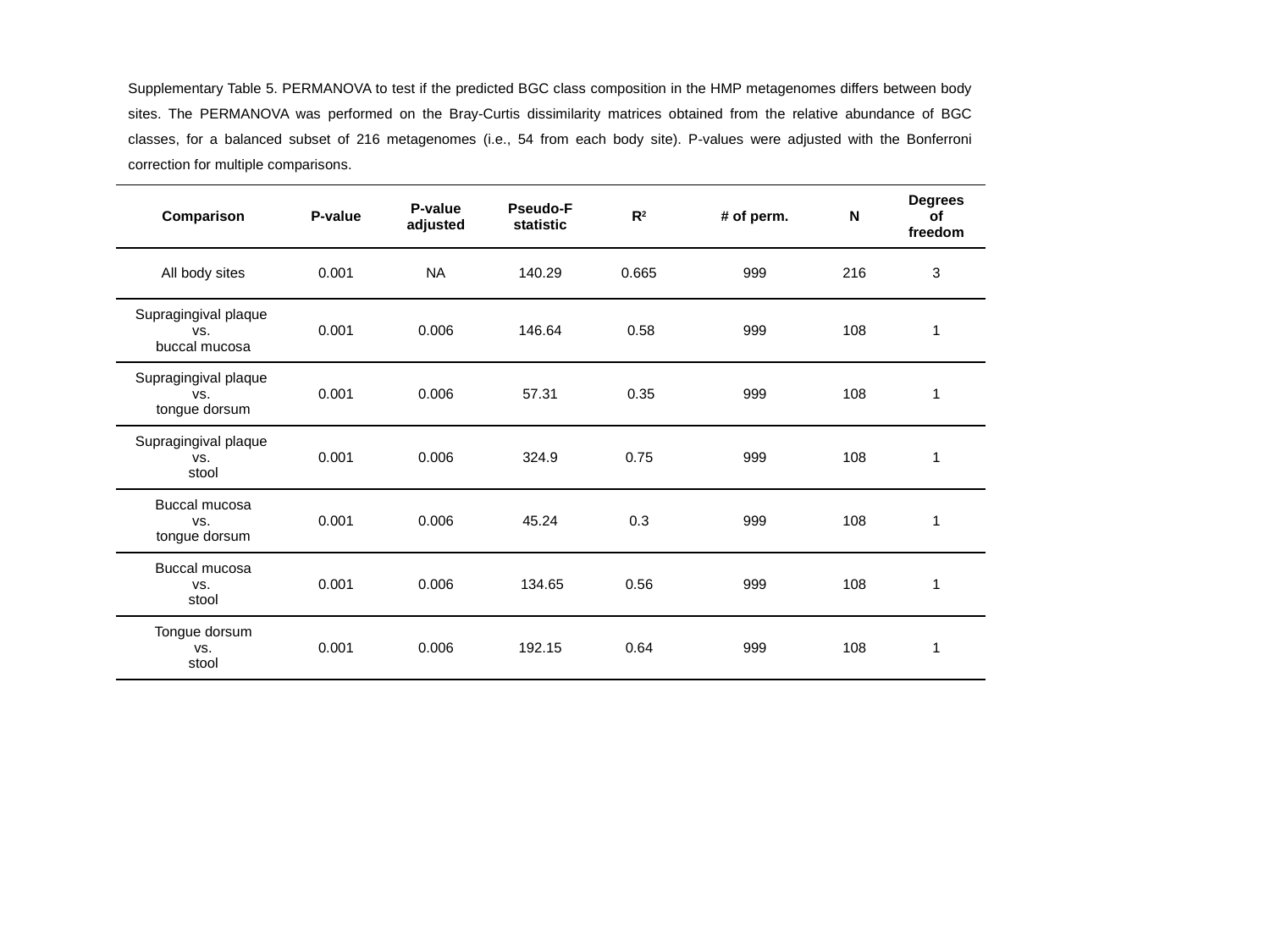

Supplementary Table 5. PERMANOVA to test if the predicted BGC class composition in the HMP metagenomes differs between body sites. The PERMANOVA was performed on the Bray-Curtis dissimilarity matrices obtained from the relative abundance of BGC classes, for a balanced subset of 216 metagenomes (i.e., 54 from each body site). P-values were adjusted with the Bonferroni correction for multiple comparisons.
| Comparison | P-value | P-value adjusted | Pseudo-F statistic | R^2^ | # of perm. | N | Degrees of freedom |
| --- | --- | --- | --- | --- | --- | --- | --- |
| All body sites | 0.001 | NA | 140.29 | 0.665 | 999 | 216 | 3 |
| Supragingival plaque vs. buccal mucosa | 0.001 | 0.006 | 146.64 | 0.58 | 999 | 108 | 1 |
| Supragingival plaque vs. tongue dorsum | 0.001 | 0.006 | 57.31 | 0.35 | 999 | 108 | 1 |
| Supragingival plaque vs. stool | 0.001 | 0.006 | 324.9 | 0.75 | 999 | 108 | 1 |
| Buccal mucosa vs. tongue dorsum | 0.001 | 0.006 | 45.24 | 0.3 | 999 | 108 | 1 |
| Buccal mucosa vs. stool | 0.001 | 0.006 | 134.65 | 0.56 | 999 | 108 | 1 |
| Tongue dorsum vs. stool | 0.001 | 0.006 | 192.15 | 0.64 | 999 | 108 | 1 |

### Slide 6
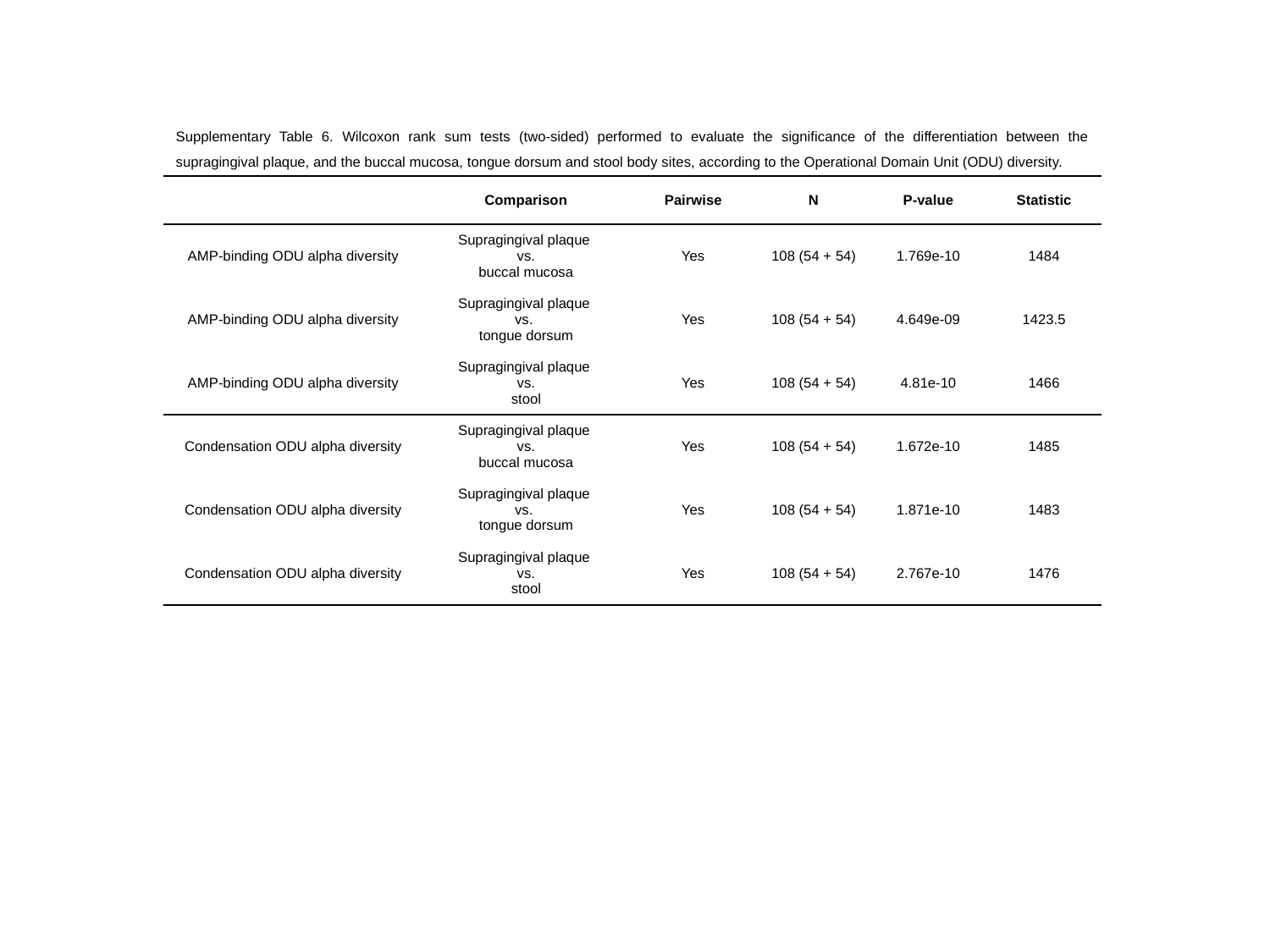

Supplementary Table 6. Wilcoxon rank sum tests (two-sided) performed to evaluate the significance of the differentiation between the supragingival plaque, and the buccal mucosa, tongue dorsum and stool body sites, according to the Operational Domain Unit (ODU) diversity.
| | Comparison | Pairwise | N | P-value | Statistic |
| --- | --- | --- | --- | --- | --- |
| AMP-binding ODU alpha diversity | Supragingival plaque vs. buccal mucosa | Yes | 108 (54 + 54) | 1.769e-10 | 1484 |
| AMP-binding ODU alpha diversity | Supragingival plaque vs. tongue dorsum | Yes | 108 (54 + 54) | 4.649e-09 | 1423.5 |
| AMP-binding ODU alpha diversity | Supragingival plaque vs. stool | Yes | 108 (54 + 54) | 4.81e-10 | 1466 |
| Condensation ODU alpha diversity | Supragingival plaque vs. buccal mucosa | Yes | 108 (54 + 54) | 1.672e-10 | 1485 |
| Condensation ODU alpha diversity | Supragingival plaque vs. tongue dorsum | Yes | 108 (54 + 54) | 1.871e-10 | 1483 |
| Condensation ODU alpha diversity | Supragingival plaque vs. stool | Yes | 108 (54 + 54) | 2.767e-10 | 1476 |

### Slide 7
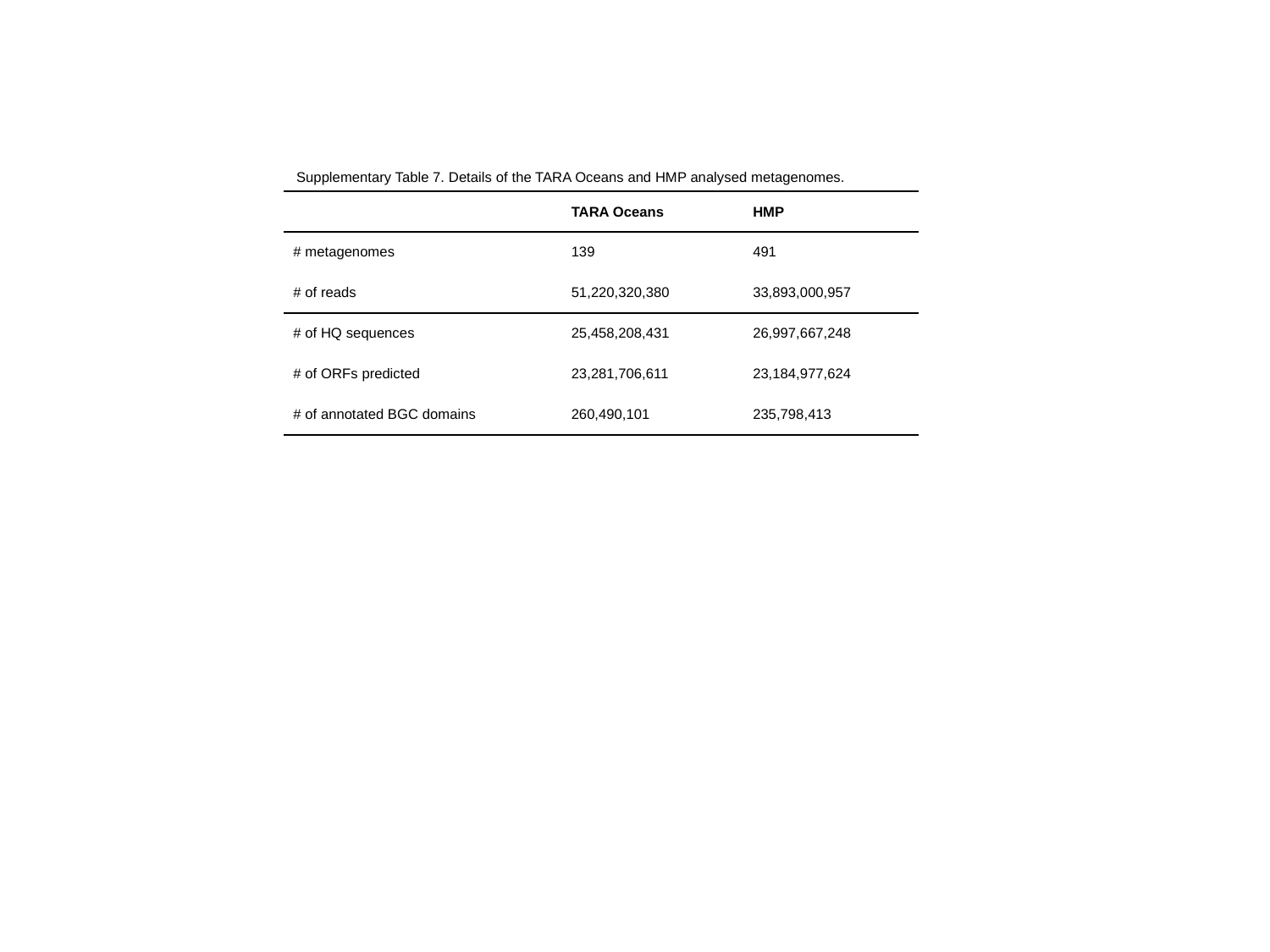

Supplementary Table 7. Details of the TARA Oceans and HMP analysed metagenomes.
| | TARA Oceans | HMP |
| --- | --- | --- |
| # metagenomes | 139 | 491 |
| # of reads | 51,220,320,380 | 33,893,000,957 |
| # of HQ sequences | 25,458,208,431 | 26,997,667,248 |
| # of ORFs predicted | 23,281,706,611 | 23,184,977,624 |
| # of annotated BGC domains | 260,490,101 | 235,798,413 |

### Slide 8
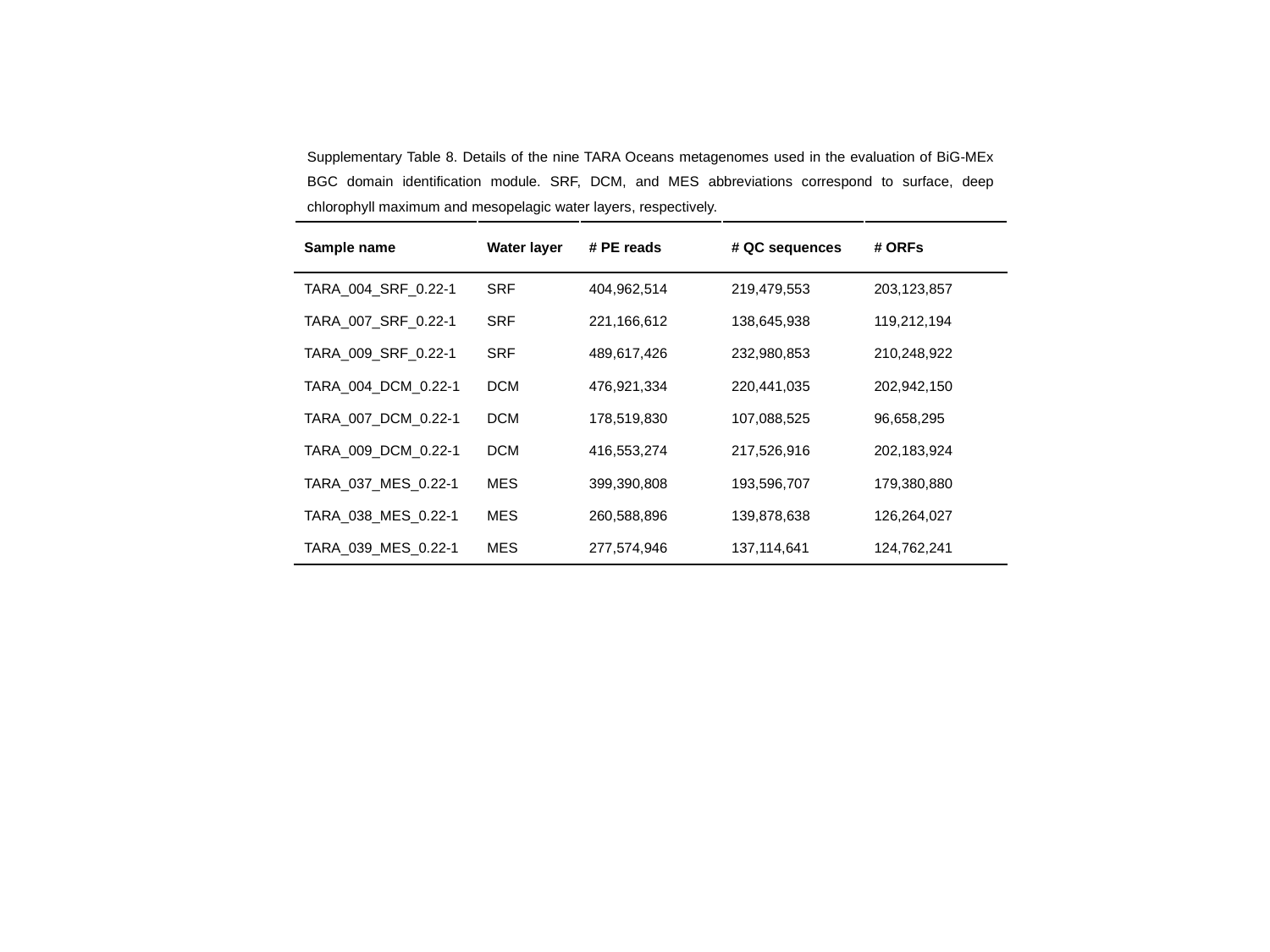

Supplementary Table 8. Details of the nine TARA Oceans metagenomes used in the evaluation of BiG-MEx BGC domain identification module. SRF, DCM, and MES abbreviations correspond to surface, deep chlorophyll maximum and mesopelagic water layers, respectively.
| Sample name | Water layer | # PE reads | # QC sequences | # ORFs |
| --- | --- | --- | --- | --- |
| TARA\_004\_SRF\_0.22-1 | SRF | 404,962,514 | 219,479,553 | 203,123,857 |
| TARA\_007\_SRF\_0.22-1 | SRF | 221,166,612 | 138,645,938 | 119,212,194 |
| TARA\_009\_SRF\_0.22-1 | SRF | 489,617,426 | 232,980,853 | 210,248,922 |
| TARA\_004\_DCM\_0.22-1 | DCM | 476,921,334 | 220,441,035 | 202,942,150 |
| TARA\_007\_DCM\_0.22-1 | DCM | 178,519,830 | 107,088,525 | 96,658,295 |
| TARA\_009\_DCM\_0.22-1 | DCM | 416,553,274 | 217,526,916 | 202,183,924 |
| TARA\_037\_MES\_0.22-1 | MES | 399,390,808 | 193,596,707 | 179,380,880 |
| TARA\_038\_MES\_0.22-1 | MES | 260,588,896 | 139,878,638 | 126,264,027 |
| TARA\_039\_MES\_0.22-1 | MES | 277,574,946 | 137,114,641 | 124,762,241 |

### Slide 9
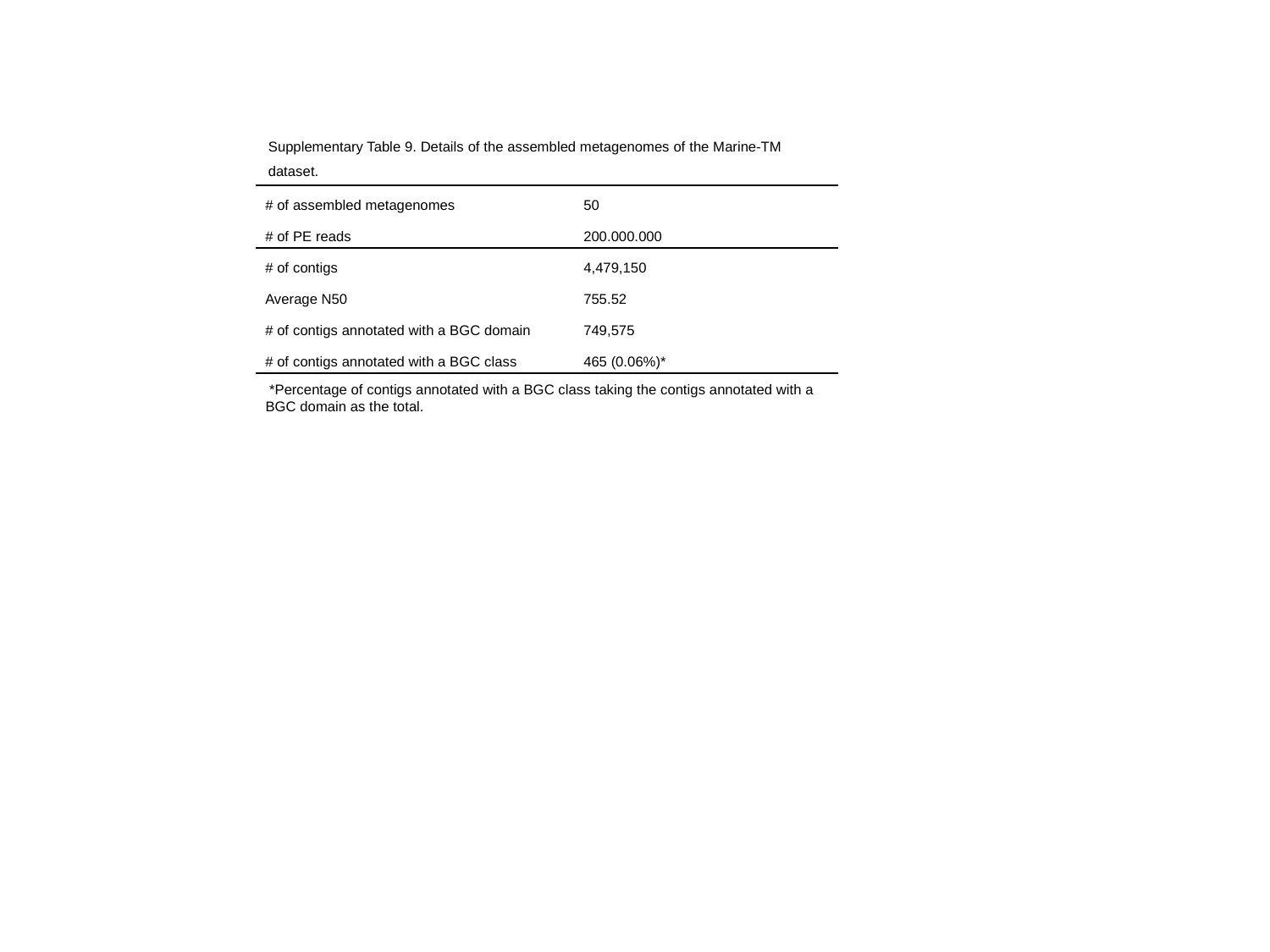

Supplementary Table 9. Details of the assembled metagenomes of the Marine-TM dataset.
| # of assembled metagenomes | 50 |
| --- | --- |
| # of PE reads | 200.000.000 |
| # of contigs | 4,479,150 |
| Average N50 | 755.52 |
| # of contigs annotated with a BGC domain | 749,575 |
| # of contigs annotated with a BGC class | 465 (0.06%)\* |
 *Percentage of contigs annotated with a BGC class taking the contigs annotated with a BGC domain as the total.
